# Metabolic independence drives gut microbial colonization and resilience in health and disease

**DOI:** 10.1101/2021.03.02.433653

**Authors:** Andrea R. Watson, Jessika Füssel, Iva Veseli, Johanna Zaal DeLongchamp, Marisela Silva, Florian Trigodet, Karen Lolans, Alon Shaiber, Emily Fogarty, Joseph M. Runde, Christopher Quince, Michael K. Yu, Arda Söylev, Hilary G. Morrison, Sonny T.M. Lee, Dina Kao, David T. Rubin, Bana Jabri, Thomas Louie, A. Murat Eren

## Abstract

Changes in microbial community composition as a function of human health and disease states have sparked remarkable interest in the human gut microbiome. However, establishing reproducible insights into the determinants of microbial succession in disease has been a formidable challenge. Here we use fecal microbiota transplantation (FMT) as an *in natura* experimental model to investigate the association between metabolic independence and resilience in stressed gut environments. Our genome-resolved metagenomics survey suggests that FMT serves as an environmental filter that favors populations with higher metabolic independence, the genomes of which encode complete metabolic modules to synthesize critical metabolites, including amino acids, nucleotides, and vitamins. Interestingly, we observe higher completion of the same biosynthetic pathways in microbes enriched in IBD patients. These observations suggest a general mechanism that underlies changes in diversity in perturbed gut environments, and reveal taxon-independent markers of ‘dysbiosis’ that may explain why widespread yet typically low abundance members of healthy gut microbiomes can dominate under inflammatory conditions without any causal association with disease.

## Introduction

Understanding the determinants of microbial colonization is one of the fundamental aims of gut microbial ecology (Costello et al. 2012; Messer et al. 2017). The gradual maturation of the microbiome during the first months of life (Stewart et al. 2018), the importance of diet and lifestyle in shaping the gut microbiome (Koenig et al. 2011; Rothschild et al. 2018), and the biogeography of microbial populations along the gastrointestinal tract (Donaldson, Lee, and Mazmanian 2016) strongly suggest the importance of niche-based interactions between the gut environment and its microbiota. Previous studies that described such interactions in the context of microbial colonization have focused on microbial succession in infant gut microbiomes (Stewart et al. 2018), or relied on model systems such as germ free mice conventionalized with a consortium of microbial isolates from infant stool (Feng et al. 2020). However, our understanding of the ecological underpinnings of secondary succession following a major ecosystem disturbance caused by complex environmental factors in the gut microbiome remains incomplete. A wide range of diseases and disorders are associated with such disturbances, (Almeida et al., 2020; Durack and Lynch, 2019; Lynch and Pedersen, 2016), however; mechanistic underpinnings of these associations have been difficult to resolve. This is in part due to the diversity of human lifestyles (David et al., 2014), and the limited utility of model systems to make robust causal inferences for microbially mediated human diseases (Walter et al., 2020).

Inflammatory bowel disease (IBD), a group of increasingly common intestinal disorders that cause inflammation of the gastrointestinal tract (Baumgart and Carding 2007), has been a model to study human diseases associated with the gut microbiota (Schirmer et al. 2019). The pathogenesis of IBD is attributed in part to the gut microbiome (Plichta et al. 2019), yet the microbial ecology of IBD-associated dysbiosis remains a puzzle. Despite marked changes in gut microbial community composition in IBD (Ott et al. 2004; Sokol and Seksik 2010; Joossens et al. 2011), the microbiota associated with the disease lacks acquired infectious pathogens (Chow, Tang, and Mazmanian 2011), and microbes that are found in IBD typically also occur in healthy individuals (Clooney et al. 2021), which complicates the search for robust functional or taxonomic markers of health and disease states (Lloyd-Price et al. 2019). One of the hallmarks of IBD is reduced microbial diversity during episodes of inflammation, when the gut environment is often dominated by microbes that typically occur in lower abundances prior to inflammation (Vineis et al. 2016). The sudden increase in the relative abundance of microbes that are also common to healthy individuals suggests that the harsh conditions of IBD likely act as an ecological filter that eliminates some populations while allowing others to bloom. Yet, in the absence of an understanding of the genetic requirements for survival in IBD, critical insights into the functional drivers of microbial community succession in such disease states remains elusive.

Fecal microbiota transplantation (FMT), the transfer of stool from a donor into a recipient’s gastrointestinal tract (Eiseman et al. 1958), represents an experimental middleground to capture complex ecological interactions that shape the microbial community during secondary succession of a disrupted gut environment. FMT is frequently employed in the treatment of recurrent *Clostridioides difficile* infection (CDI) (van Nood et al. 2013) that can cause severe diarrhea and intestinal inflammation. In addition to its medical utility, FMT offers a powerful framework to study fundamental questions of microbial ecology by colliding the microbiome of a healthy donor with the disrupted gut environment of the recipient. The process presents an ecological filter with the potential to reveal functional determinants of microbial colonization success and resilience in impaired gut environments (Schmidt, Raes, and Bork 2018).

Here we use FMT as an *in natura* experimental model to investigate the ecological and functional determinants of successful colonization of the human gut at the level of individual microbial populations using genome-resolved metagenomics. Our findings highlight the importance of environmental selection acting on the biosynthetic capacity for essential nutrients as a key driver of colonization outcome after FMT and resilience during inflammation, and demonstrate that metabolic independence serves as a taxonomy-independent determinant of colonization success in the human gut.

## Results and Discussion

### Study Design

Our study includes 109 gut metagenomes (Supplementary Table 1) from two healthy FMT donors (A and B) and 10 FMT recipients (five recipients per donor) with multiple recurrent CDI. We collected 24 Donor A samples over a period of 636 days and 15 Donor B samples over a period of 532 days to establish an understanding of the long-term microbial population dynamics within each donor microbiota. The FMT recipients received vancomycin for a minimum of 10 days to attain resolution of diarrheal illness prior to FMT. On the last day of vancomycin treatment, a baseline fecal sample was collected from each recipient, and their bowel contents were evacuated immediately prior to FMT. Recipients did not take any antibiotics on the day of transplant, or during the post-FMT sampling period (Supplementary Figure 1). We collected 5 to 9 samples from each recipient for a period of up to 336 days post-FMT. Deep sequencing of donor and recipient metagenomes using Illumina paired-end (2×150) technology resulted in a total of 7.7 billion sequences with an average of 71 million reads per metagenome (Figure 1, Supplementary Table 1, Supplementary Table 2). We employed genome-resolved metagenomics, microbial population genetics, and metabolic pathway reconstruction for an in-depth characterization of donor and recipient gut microbiotas, and we leveraged publicly available gut metagenomes to benchmark our observations.

**Figure 1.**
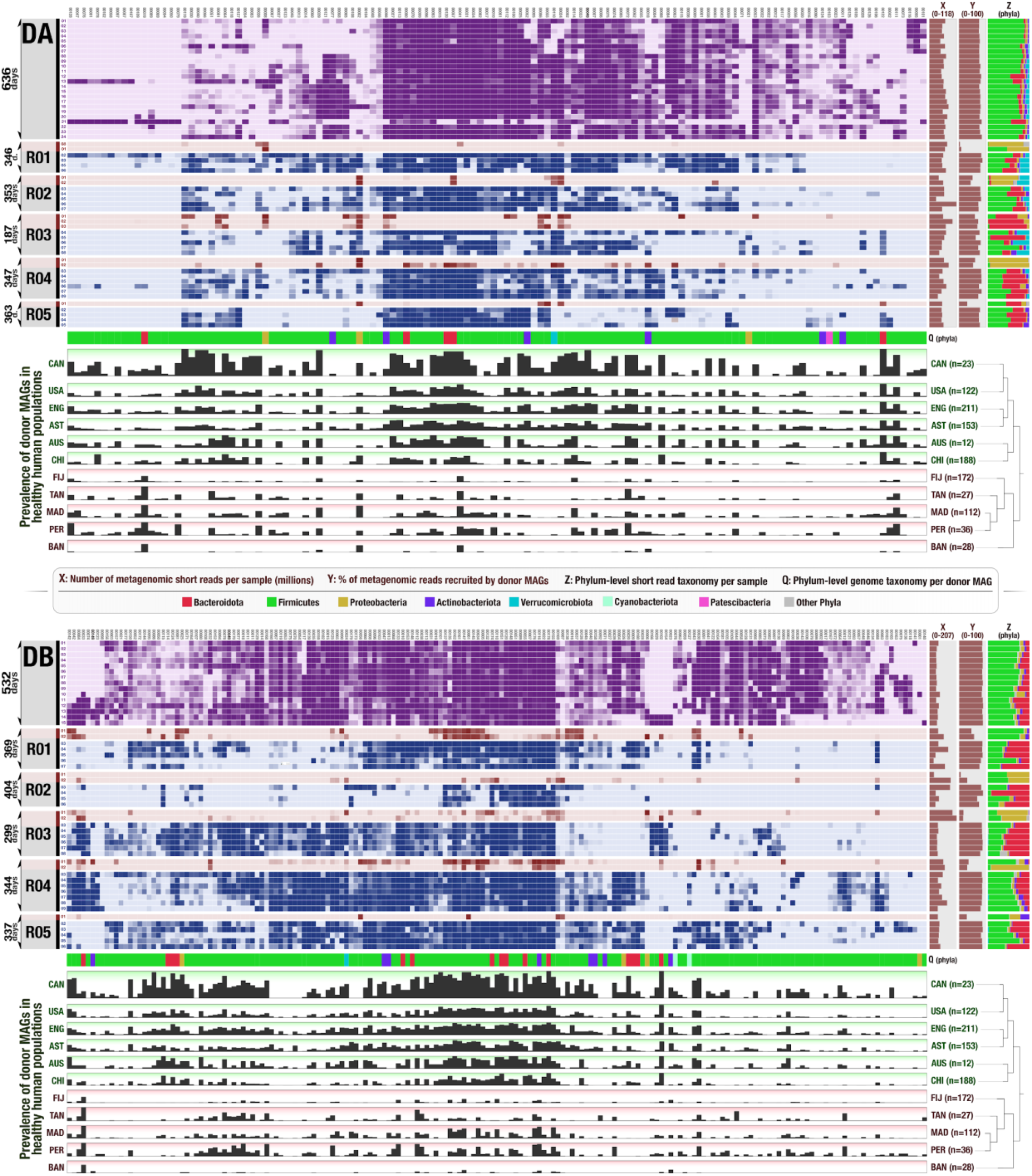
Detection of FMT donor genomes in FMT recipients and publicly available gut metagenomes. In both heat maps each column represents a donor genome, each row represents a metagenome, and each data point represents the detection of a given genome in a given metagenome. Genomes are clustered according to their detection in all metagenomes (Euclidean distance and Ward clustering). Purple rows represent donor metagenomes from stool samples collected over 636 days for Donor A and 532 days for Donor B. Red rows represent recipient pre-FMT metagenomes, and blue rows represent recipient post-FMT metagenomes. The three rightmost columns display for each metagenome: (X) the number of metagenomic short reads in millions, (Y) the percent of metagenomic short reads recruited by genomes, and (Z) the taxonomic composition of metagenomes (based on metagenomic short reads) at the phylum level. The first row below each heat map (Q) provides the phylum-level taxonomy for each donor genome. Finally, the 11 bottommost rows under each heat map show the fraction of healthy adult metagenomes from 11 different countries in which a given donor genome is detected (if a genome is detected in every individual from a country it is represented with a full bar). The dendrograms on the right-hand side of these layers organize countries based on the detection patterns of genomes (Euclidean distance and Ward clustering). Red and green shades represent the two main clusters that emerge from this analysis, where green layers are industrialized countries in which donor genomes are highly prevalent and red layers are less industrialized countries where the prevalence of donor genomes is low. A high resolution version of this figure is also available at https://doi.org/10.6084/m9.figshare.15138720.

### Genome-resolved metagenomics show many, but not all, donor microbes colonized recipients and persisted long-term

We first characterized the taxonomic composition of each donor and recipient sample by analyzing our metagenomic short reads given a clade-specific k-mer database (Supplementary Table 2). The phylum-level microbial community composition of both donors reflected those observed in healthy individuals in North America (Human Microbiome Project Consortium 2012): a large representation of Firmicutes and Bacteroidetes, and other taxa with lower relative abundances including Actinobacteria, Verrucomicrobia, and Proteobacteria (Figure 1, Supplementary Table 2). In contrast, the vast majority of the recipient pre-FMT samples were dominated by Proteobacteria, a phylum that typically undergoes a drastic expansion in individuals treated with vancomycin (Isaac et al. 2017). After FMT, we observed a dramatic shift in recipient taxonomic profiles (Supplementary Table 2, Supplementary Figure 2, Supplementary Figure 3), a widely documented hallmark of this procedure (Khoruts et al. 2010; Grehan et al. 2010; Shahinas et al. 2012). Nearly all recipient samples post-FMT were dominated by Bacteroidetes and Firmicutes as well as Actinobacteria and Verrucomicrobia in lower abundances, resembling qualitatively, but not quantitatively, the taxonomic profiles of their donors (Supplementary Table 2). The phylum Bacteroidetes was over-represented in recipients: even though the median relative abundance of Bacteroidetes populations were 5% and 17% in donors A and B, their relative abundance in recipients post-FMT was 33% and 45%, respectively (Figure 1, Supplementary Table 2). A single genus, *Bacteroides*, made up 76% and 82% of the Bacteroidetes populations in the recipients of Donor A and B, respectively (Supplementary Table 2). The success of the donor *Bacteroides* populations in recipients upon FMT is not surprising given the ubiquity of this genus across geographically diverse human populations (Wexler and Goodman 2017) and the ability of its members to survive substantial levels of stress (Swidsinski et al. 2005; Vineis et al. 2016). This initial coarse taxonomic analysis demonstrates the successful transfer of only some populations, suggesting selective filtering of the transferred community.

To generate insights into the genomic content of the microbial community, we first assembled short metagenomic reads into contiguous segments of DNA (contigs). Co-assemblies of 24 Donor A and 15 Donor B metagenomes independently resulted in 53,891 and 54,311 contigs that were longer than 2,500 nucleotides and described 0.70 and 0.79 million genes occurring in 179 and 248 genomes, as estimated by the mode of the frequency of bacterial single-copy core genes (Supplementary Table 2). On average, 80.8% of the reads in donor metagenomes mapped back to the assembled contigs from donor metagenomes, which suggests that the assemblies represented a large fraction of the donor microbial communities. Donor assemblies recruited only 43.4% of the reads on average from the pre-FMT recipient metagenomes. This number increased to 80.2% for post-FMT recipient metagenomes and remained at an average of 76.8% even one year post-FMT (Supplementary Table 2). These results suggest that members of the donor microbiota successfully established in the recipient gut and persisted long-term.

To investigate functional determinants of microbial colonization by identifying donor populations that were successful at colonizing multiple individuals, we reconstructed microbial genomes from donor assemblies using sequence composition and differential coverage signal as previously described (Sharon et al. 2013; Lee et al. 2017). We manually refined metagenomic bins to improve their quality following previously described approaches (Delmont et al. 2018; Shaiber et al. 2020) and only retained those that were at least 70% complete and had no more than 10% redundancy as predicted by bacterial single-copy core genes (Bowers et al. 2017; Chen et al. 2020). Our binning effort resulted in a final list of 128 metagenome-assembled genomes (MAGs) for Donor A and 183 MAGs for Donor B that included members of Firmicutes (n=265), Bacteroidetes (n=20), Actinobacteria (n=14), Proteobacteria (n=7), Verrucomicrobia (n=2), Cyanobacteria (n=2), and Patescibacteria (n=1) (Supplementary Table 3). The taxonomy of donor-derived genomes largely reflected the taxonomic composition of donor metagenomic short reads (Figure 1, Supplementary Table 2, Supplementary Table 3). While only 20 genomes (mostly of the genera *Bacteroides* and *Alistipes*) explained the entirety of the Bacteroidetes group, we recovered 265 genomes that represented lower abundance but diverse populations of Firmicutes (Figure 1, Supplementary Table 2, Supplementary Table 3).

### Metagenomic read recruitment elucidates colonization events

Reconstructing donor genomes enabled us to characterize (1) population-level microbial colonization dynamics before and after FMT using donor and recipient metagenomes and (2) the distribution of each donor population across geographically distributed humans using 1,984 publicly available human gut metagenomes (Figure 1, Supplementary Table 4).

Our metagenomic read recruitment analysis showed that donor A and B genomes recruited on average 77.05% and 83.04%, respectively, of reads from post-FMT metagenomes, suggesting that the collection of donor genomes well represents the recipient metagenomes post-FMT (Figure 1). As expected, we detected each donor population in at least one donor metagenome (see Methods for ‘detection’ criteria). Yet, only 16% of Donor A populations were detected in every Donor A sample, and only 44% of Donor B populations were detected in every Donor B sample (Figure 1, Supplementary Table 3), demonstrating the previously documented dynamism of gut microbial community composition over time (David et al. 2014). A marked increase in the detection of donor populations in recipients after FMT is in agreement with the general pattern of transfer suggested by the short-read taxonomy (Figure 1): while we detected only 38% of Donor A and 54% of Donor B populations in at least one recipient pre-FMT, these percentages increased to 96% for both donors post-FMT (Supplementary Table 3). We note that we observed a higher fraction of donor populations in recipients as a function of the FMT delivery method. Following the cases of FMT where donor stool was transplanted via colonoscopy, we detected 54.7% and 33.3% donor genomes in the recipients of donor A (n=3) and donor B (n=2), respectively. In contrast, in the cases of FMT where donor stool was transplanted via pills, we detected 69.5% and 61.6% donor genomes in the recipients of donor A (n=2) and donor B (n=3), respectively.

Not every donor population was detected in each recipient, but the emergence of donor populations in recipients did not appear to be random: while some donor populations colonized all recipients, others colonized none (Figure 1), providing us with an opportunity to quantify colonization success for each donor population in our dataset.

### Succession of donor microbial populations in FMT recipients and their prevalence in publicly available metagenomes reveal good and poor colonizers

Of the populations that consistently occurred in donor metagenomes, some were absent in all or most recipient metagenomes after FMT, and others were continuously present throughout the sampling period in both donor and recipient metagenomes (Figure 1). To gain insights into the ecology of donor microbial populations beyond our dataset, we explored their occurrence in publicly available healthy gut metagenomes through metagenomic read recruitment. This analysis enabled us to consider the prevalence of donor populations in FMT recipients and global gut metagenomes and to define two groups of donor genomes that represented opposite colonization and prevalence phenotypes.

The ‘good colonizers’ comprise those microbial populations that colonized and persisted in all FMT recipients. Intriguingly, these populations were also the most prevalent in publicly available gut metagenomes from Canada. Overall, these donor microbial populations (1) systematically colonized the majority of FMT recipients, (2) persisted in these environments long-term regardless of host genetics or lifestyle, and (3) were prevalent in public gut metagenomes outside of our study. In contrast, the so-called ‘poor colonizers’ failed to colonize or persist in at least three FMT recipients. These populations were nevertheless viable in the donor gut environment: not only did they occur systematically in donor metagenomes, but they also sporadically colonized some FMT recipients. Yet, unlike the good colonizers, the distribution patterns of poor colonizers were sparse within our cohort, as well as within the publicly available metagenomes. In fact, populations identified as poor colonizers were less prevalent than good colonizers in each of the 17 different countries we queried. In countries including the United States, Canada, Austria, China, England, and Australia, microbial populations identified as good colonizers occurred in 5 times more people than poor colonizers in the same country (Figure 1, Supplementary Table 3), which suggests that the outcomes of FMT in our dataset were unlikely determined by neutral processes. This observation is in contrast with previous studies that suggested ‘dose’ (i.e., the abundance of a given population in donor fecal matter) as a predominant force that determines outcomes of colonization after FMT (Podlesny and Florian Fricke, 2020; Smillie et al., 2018). However, our strain-resolved analysis of colonization events in our data in conjunction with the distribution of the same populations in publicly available metagenomes revealed (1) a significant correlation between the colonization success of donor populations and their prevalence across publicly available metagenomes, and (2) showed that the prevalence of a given population across global gut metagenomes can predict its colonization success after FMT better than its abundance in the donor stool sample (Wald test, p=6.3e-06 and p=9.0e-07) (Supplementary Information). Overall, these observations suggest a link between the colonization outcomes in our study and global prevalence of the same microbial populations, and that the succession of donor populations in our data were likely influenced by selective processes that influence colonization outcomes.

Next, we sought to investigate whether we can identify metabolic features that systematically differ between good colonizers and poor colonizers independent of their taxonomy. To conduct such a comparative analysis, we conservatively selected the top 20 populations from each group that best reflect their group properties by considering both their success after FMT and their prevalence across publicly available metagenomes (Supplementary Table 7). The 20 populations representative of good colonizers were dominated by Firmicutes (15 of 20) but also included Bacteroidetes and one Actinobacteria population. All populations identified as poor colonizers resolved to Firmicutes (Figure 2, Supplementary Table 7). Genome completion estimates did not differ between good and poor colonizers (Wilcoxon rank sum test, p=0.42) and averaged to 91% and 93%, respectively. But intriguingly, the genome sizes between the two groups differed dramatically (p=2.9e-06): genomes of good colonizers averaged to 2.8 Mbp while those of poor colonizers averaged to 1.6 Mbp. We considered that our bioinformatics analyses may have introduced biases to genome lengths, but found a very high correspondence between the lengths of the genomes and their best matching reference genomes in the Genome Taxonomy Database (GTDB) (R^2^=0.88, p=5e-14). Assuming that the generally larger genomes of good colonizers may be an indication of an increased repertoire of core metabolic competencies compared to poor colonizers, we next conducted a metabolic enrichment analysis for quantitative insights (see Materials and Methods).

**Figure 2.**
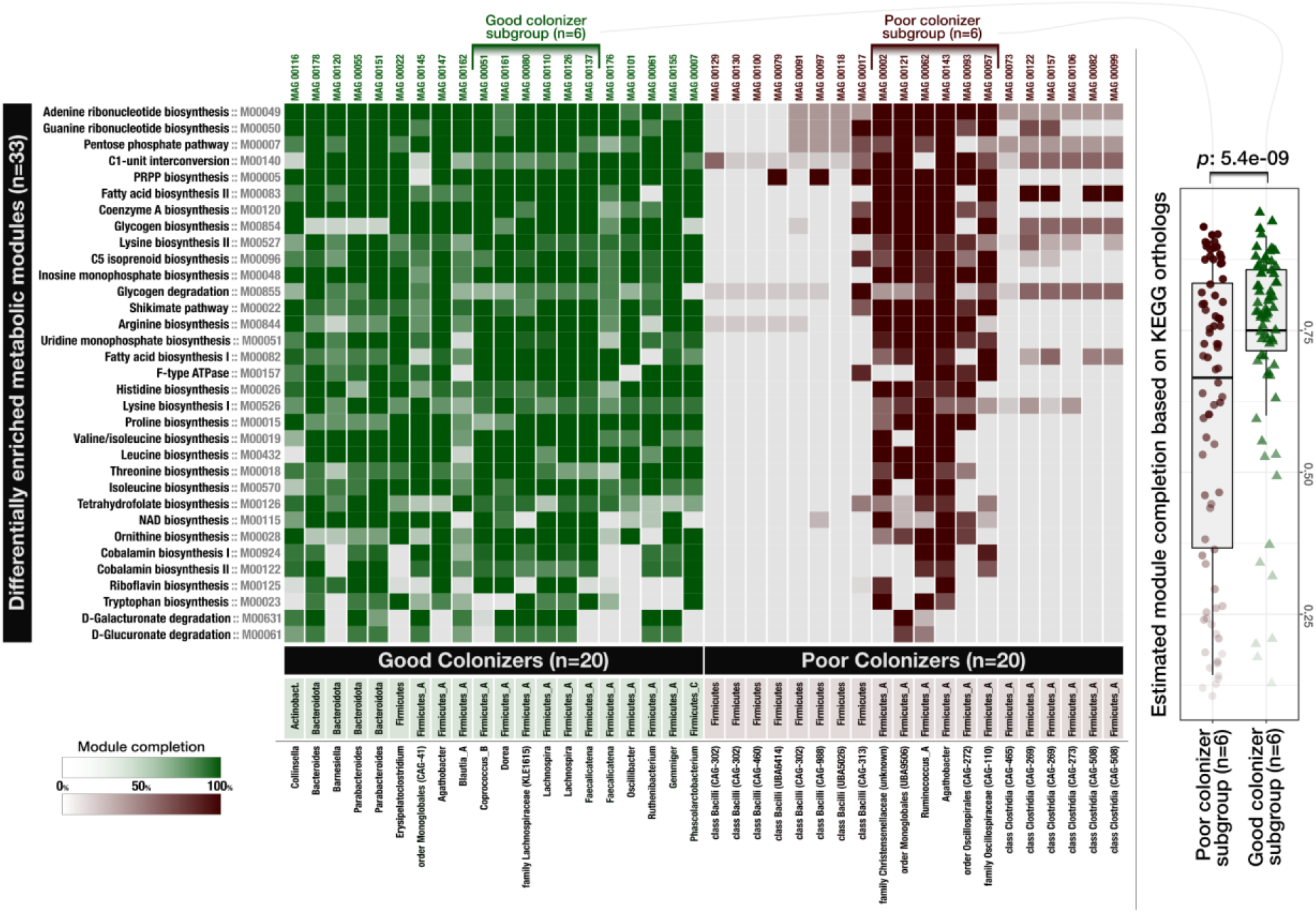
Distribution of metabolic modules across genomes of good and poor colonizers. Each data point in this heat map shows the level of completion of a given metabolic module (rows) in a given genome (columns). The box-plot on the right-side compares a subset of poor colonizer and good colonizer genomes, where each data point represents the level of completion of a given metabolic module in a genome and shows a statistically significant difference between the overall completion of metabolic modules between these subgroups (Wilcoxon rank sum test, p=5.4e-09). A high-resolution version of this figure is also available at https://doi.org/10.6084/m9.figshare.15138720.

### Good colonizers are enriched in metabolic pathways for the biosynthesis of essential organic compounds

Our enrichment analysis between good and poor colonizers revealed 33 metabolic modules that were enriched in good colonizers and none that were enriched in poor colonizers (Figure 2, Supplementary Table 7). Of all enriched modules, 79% were related to biosynthesis, indicating an overrepresentation of biosynthetic capabilities among good colonizers as KEGG modules for biosynthesis only make up 55% of all KEGG modules (Figure 2, Supplementary Table 7). Of the 33 enriched modules, 48.5% were associated with amino acid metabolism, 21.2% with vitamin and cofactor metabolism, 18.2% with carbohydrate metabolism, 24.2% with nucleotide metabolism, 6% with lipid metabolism and 3% with energy metabolism (Supplementary Table 7). Metabolic modules that were enriched in the good colonizers included the biosynthesis of seven of nine essential amino acids, indicating the importance of high metabolic independence to synthesize essential compounds as a likely factor that increases success in colonizing new environments (Supplementary Table 7). This is further supported by the enrichment of biosynthesis pathways for the essential cofactor vitamin B12 (cobalamin), which occurred in 67.5% of the good colonizers and only 12.5% of the poor colonizers (Supplementary Table 7). Vitamin B12 is structurally highly complex and costly to produce, requiring expression of more than 30 genes that are exclusively encoded by bacteria and archaea (J. H. Martens et al. 2002). In addition to the biosynthesis of tetrahydrofolate, riboflavin, and cobalamin, the genomes of good colonizers had a larger representation of biosynthetic modules for vitamins including biotin, pantothenate, folate, and thiamine (Supplementary Table 7). These micronutrients are equally essential in bacterial and human metabolism and are important mediators of host-microbe interactions (Biesalski 2016). Interestingly, enriched metabolic modules in our analysis partially overlap with those that Feng *et al*. identified as the determinants of microbial fitness using metatranscriptomics and a germ-free mouse model conventionalized with microbial isolates of human origin (Feng et al. 2020).

Even though these 33 metabolic modules were statistically enriched in populations identified as good colonizers, some of them also occurred in the genomes of poor colonizers (Figure 2). To identify whether the levels of completion of these modules could distinguish the good and poor colonizers, we matched six good colonizers that encoded modules enriched in these populations to six populations of poor colonizers from the same phylum (Figure 2). Bacterial single-copy core genes estimated that genomes in both subgroups were highly complete with a slight increase in average genome completion of poor colonizers (93.7%) compared to good colonizers (90.1%). Despite the higher estimated genome completion for populations of poor colonizers, estimated metabolic module completion values were slightly yet significantly lower in this group (Wilcoxon rank sum test with continuity correction, V=958, p=5e-09) (Figure 2, Supplementary Table 7). Thus, these modules were systematically missing genes in populations of poor colonizers, indicating their functionality was likely reduced, if not absent.

These observations suggest that the ability to synthesize cellular building blocks, cofactors and vitamins required for cellular maintenance and growth provides a substantial advantage during secondary succession, highlighting that the competitive advantages conferred by metabolic autonomy may outweigh the additional costs under certain conditions. For the remainder of our study, we use the term ‘high metabolic independence’ (HMI) to describe genomic evidence for a population’s ability to synthesize essential compounds, and ‘low metabolic independence’ (LMI) to describe the absence of, or reduction in, such capacity.

### While gut microbial ecosystems of healthy individuals include microbes with both low- and high-metabolic independence, IBD primarily selects for microbes with high-metabolic independence

Our results so far show that while the healthy donor environment could support both HMI and LMI populations (Figure 1, Supplementary Table 3), challenging microbes to colonize a new environment or to withstand ecosystem perturbation during FMT selects for HMI populations (Figure 2, Supplementary Table 7), suggesting that metabolic independence is a more critical determinant of fitness during stress than during homeostasis. Based on these observations, it is conceivable to hypothesize that (1) a gut environment in homeostasis will support a large variety of microbial populations with a wide spectrum of metabolic independence, and (2) a gut environment under stress will select for populations with high metabolic independence, potentially leading to an overall reduction in diversity.

To test these hypotheses, we compared genomes reconstructed from a cohort of healthy individuals (Pasolli et al. 2019) to genomes reconstructed from individuals who were diagnosed with inflammatory bowel disease (IBD). Our IBD dataset was composed of two cohorts: a set of patients with pouchitis (Vineis et al. 2016), a form of IBD with similar pathology to ulcerative colitis (De Preter et al. 2009), and a set of pediatric Crohn’s disease patients (Quince et al. 2015). The number of genomes per individual and the average level of genome completeness per group were similar between healthy individuals and those with IBD: overall, our analysis compared 264 genomes from 22 healthy individuals with an average completion of 90.4%, 44 genomes from 4 pouchitis patients with an average completion of 89.2% and 256 genomes from 12 Crohn’s disease patients with an average completion of 94.1% (Supplementary Table 8). Intriguingly, similar to the size differences between genomes of HMI populations and LMI populations (2.8 Mbp versus 1.6 Mbp on average), genomes of microbial populations associated with IBD patients were larger compared to those of microbial populations in healthy people and averaged to 3.0 Mbp versus 2.6 Mbp, respectively (Supplementary Table 8). This suggests that the environmental filters created by FMT and gastrointestinal inflammation both select for microbial populations with larger genomes and potentially higher metabolic independence.

Next, we asked whether the completion of metabolic modules associated with colonization success and resilience during FMT differed between the genomes reconstructed from healthy and IBD individuals. The completion of the 33 metabolic modules was almost identical between the HMI populations revealed by FMT and microbial populations in IBD patients (Wilcoxon rank sum test, p=0.5) (Figure 3, Supplementary Table 8). In contrast, the completion of these metabolic modules was significantly reduced in microbial populations in healthy individuals (Wilcoxon rank sum test, p < 1e-07) (Figure 3, Supplementary Table 8). Metabolic modules with the largest differences in completion between genomes from healthy and IBD individuals included biosynthesis of cobalamin, arginine, ornithine, tryptophan, isoleucine as well as the Shikimate pathway (Figure 3, Supplementary Table 8), a seven step metabolic route bacteria use for the biosynthesis of aromatic amino acids (phenylalanine, tyrosine, and tryptophan) (Herrmann and Weaver 1999).

**Figure 3.**
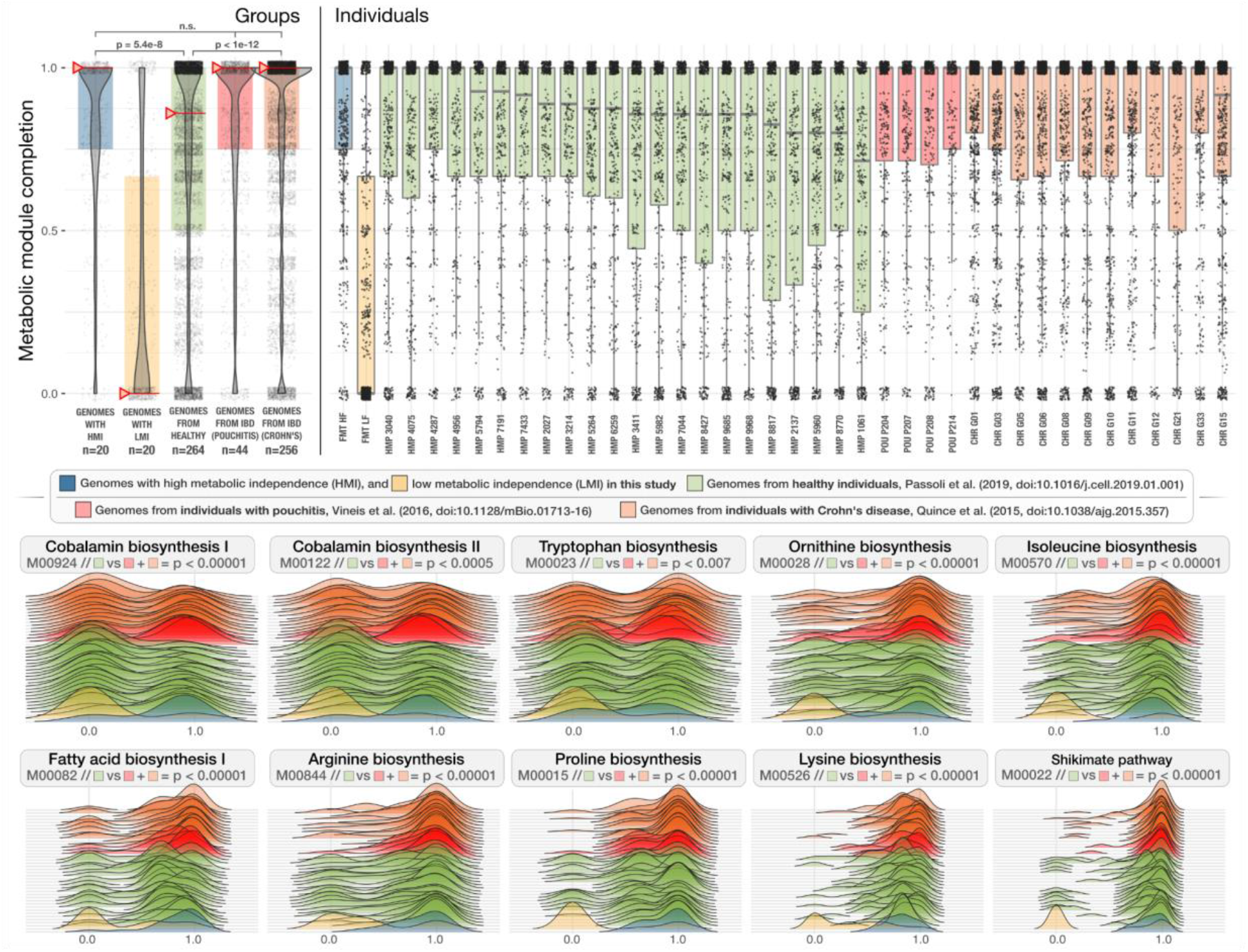
Distribution of metabolic modules in genomes reconstructed from healthy individuals and individuals with IBD. The boxplots in the top panels show the metabolic module completion values for (1) high- and (2) low-metabolic independence donor genomes identified in this study (blue and yellow), (3) genomes from healthy individuals (green), and (4) genomes from individuals with pouchitis (red) and Crohn’s disease (orange). Each dot in a given box-plot represents one of 33 metabolic modules that were enriched in HMI FMT donor populations and the y-axis indicates its estimated completion. The leftmost top panel represents group averages and red whiskers indicate the median. The rightmost top panel shows the distribution of metabolic modules for individuals within each group. In the bottom panel the completion values for 10 of the 33 pathways are demonstrated as ridge-line plots. Each plot represents a single metabolic module where each layer corresponds to an individual, and the shape of the layer represents the completion of a given metabolic module across all genomes reconstructed from that individual. A high-resolution version of this figure is also available at https://doi.org/10.6084/m9.figshare.15138720.

Our findings show that the same set of biosynthetic metabolic modules that distinguish good and poor colonizers during FMT were also differentially associated with populations of IBD patients and healthy individuals. In particular, while healthy individuals harbored microbes with a broad spectrum of metabolic capacity, microbes from individuals who suffer from two different forms of IBD had significantly higher biosynthetic independence. It is conceivable that a stable gut microbial ecosystem is more likely to support LMI populations through metabolic cross-feeding, where vitamins, amino acids, and nucleotides are exchanged between microbes (D’Souza et al. 2018). In contrast, host-mediated environmental stress in IBD likely disrupts such interactions and creates an ecological filter that selects for metabolic independence, which subsequently leads to loss of diversity and the dominance of organisms with large genomes that are often not as abundant or as competitive in states of homeostasis.

These observations have implications for our understanding of the hallmarks of healthy gut microbial ecosystems. Defining the ‘healthy gut microbiome’ has been a major goal of human gut microbiome research (Bäckhed et al. 2012), which still remains elusive (Eisenstein 2020). Despite comprehensive investigations that considered core microbial taxa (Arumugam et al. 2011; Lloyd-Price, Abu-Ali, and Huttenhower 2016) or guilds of microbes that represent coherent functional groups (Wu et al. 2021), the search for ‘biomarkers’ of healthy gut microbiomes is ongoing (McBurney et al. 2019). Our findings indicate that beyond the taxonomic diversity of a microbial community, a broad range of metabolic independence represents a defining feature of a healthy gut microbiome. Conversely, our findings also suggest that an enrichment of metabolically independent populations could serve as an indicator of environmental stress in the human gut. Detection of these metabolic markers is not influenced by fluctuations in taxonomic composition or diversity, and represents a quantifiable feature of microbial communities through genome-resolved metagenomic surveys.

Our findings offer a new, taxonomy-independent perspective on the determinants of microbial resilience in the human gut environment under stress. Yet, our study is limited to well-known metabolic pathways, which, given the extent of the unknown coding space in microbial genomes (Vanni et al. 2020), are likely far from complete. Thus, the enrichment of biosynthetic modules in HMI populations suggests that the ability to synthesize essential biological compounds is necessary but likely insufficient to survive environmental stress in the gut. Nevertheless, the finding that the same metabolic modules that promote colonization success after FMT are also the hallmarks of resilience in IBD suggests the presence of unifying ecological principles that govern microbial diversity in distinct modes of stress, which warrants deeper investigation. Research into the development of microbial therapeutics to reinstate homeostasis is continuously intensifying (Jimenez et al., 2019) and the benefits of such treatments will critically depend on successful engraftment of individual populations. Thus, considering their degree of metabolic independence may prove crucial.

## Conclusions

Our study identifies high metabolic independence conferred by the biosynthetic capacity for amino acids, nucleotides, and essential micronutrients as a distinguishing hallmark of microbial populations that colonize recipients of FMT and that thrive in IBD patients. These findings highlight the functional complexity of the human gut microbiome whose various interactions with the host are shaped through a network of microbial interactions such as cross-feeding of macro- and micro-nutrients. Our study offers a simple model that posits the following: microbial populations that are metabolically independent and those that lack the means to synthesize essential metabolites co-occur in a healthy gut environment in harmony, where their differential resilience to stress is indiscernible by their taxonomy or relative abundance. However, the challenges associated with the transfer to a new gut environment through FMT, or with host-mediated stress through IBD, initiate an ecological filter that selects for microbes that can self-sustain in the absence of ecosystem services associated with states of homeostasis. This model provides a hypothesis that explains the dominance of low-abundance members of healthy gut environments under stressful conditions, without any necessary direct causal association with disease state. If the association between particular microbial taxa and disease is solely driven by their superior metabolic independence, microbial therapies that aim to treat complex diseases by adding microbes associated with healthy individuals will be unlikely to compete with the adaptive processes that regulate complex gut microbial ecosystems.

## Materials and Methods

### Sample collection and storage

We used a subset of individuals who participated in a randomized clinical trial (Kao et al. 2017) and conducted a longitudinal FMT study of two human cohorts (DA and DB), each consisting of one FMT donor and 5 FMT recipients of that donor’s stool. All recipients received vancomycin for a minimum of 10 days pre-FMT at a dose of 125 mg four times daily. Three DA and two DB recipients received FMT via pill, and two DA and three DB recipients received FMT via colonoscopy. All recipients had recurrent *C. difficile* infection before FMT, and two DA recipients and one DB recipient were also diagnosed with ulcerative colitis (UC). 24 stool samples were collected from the DA donor over a period of 636 days, and 15 stool samples were collected from the DB donor over a period of 532 days. Between 5 and 9 stool samples were collected from each recipient over periods of 187 to 404 days, with at least one sample collected pre-FMT and 4 samples collected post-FMT. This gave us a total of 109 stool samples from all donors and recipients. Samples were stored at −80°C. (Supplementary Figure 1, Supplementary Table 1)

### Metagenomic short-read sequencing

We extracted the genomic DNA from frozen samples according to the centrifugation protocol outlined in MoBio PowerSoil kit with the following modifications: cell lysis was performed using a GenoGrinder to physically lyse the samples in the MoBio Bead Plates and Solution (5–10 min). After final precipitation, the DNA samples were resuspended in TE buffer and stored at −20 °C until further analysis. Sample DNA concentrations were determined by PicoGreen assay. DNA was sheared to ∼400 bp using the Covaris S2 acoustic platform and libraries were constructed using the Nugen Ovation Ultralow kit. The products were visualized on an Agilent Tapestation 4200 and size-selected using BluePippin (Sage Biosciences). The final library pool was quantified with the Kapa Biosystems qPCR protocol and sequenced on the Illumina NextSeq500 in a 2 × 150 paired-end sequencing run using dedicated read indexing.

### ‘Omics workflows

Whenever applicable, we automated and scaled our ‘omics analyses using the bioinformatics workflows implemented by the program ‘anvi-run-workflow’ (Shaiber et al. 2020) in anvi’o 7.1 (Eren et al. 2015, 2021). Anvi’o workflows implement numerous steps of bioinformatics tasks including short-read quality filtering, assembly, gene calling, functional annotation, hidden Markov model search, metagenomic read-recruitment, metagenomic binning, and phylogenomics. Workflows use Snakemake (Köster and Rahmann 2012) and a tutorial is available at the URL http://merenlab.org/anvio-workflows/. The following sections detail these steps.

### Taxonomic composition of metagenomes based on short reads

We used Kraken2 v2.0.8-beta (Wood, Lu, and Langmead 2019) with the NCBI’s RefSeq bacterial, archaeal, viral and viral neighbors genome databases to calculate the taxonomic composition within short-read metagenomes.

### Assembly of metagenomic short reads

To minimize the impact of random sequencing errors in our downstream analyses, we used the program ‘iu-filter-quality-minoche’ to process short metagenomic reads, which is implemented in illumina-utils v2.11 (Eren et al. 2013) and removes low-quality reads according to the criteria outlined by Minoche et al. (Minoche, Dohm, and Himmelbauer 2011). IDBA_UD v1.1.2 (Peng et al. 2012) assembled quality-filtered short reads into longer contiguous sequences (contigs), although we needed to recompile IDBA_UD with a modified header file so it could process 150bp paired-end reads.

### Processing of contigs

We use the following strategies to process both sequences we obtained from our assemblies and those we obtained from reference genomes. Briefly, we used (1) ‘anvi-gen-contigs-database’ on contigs to compute k-mer frequencies and identify open reading frames (ORFs) using Prodigal v2.6.3 (Hyatt et al. 2010), (2) ‘anvi-run-hmms’ to identify sets of bacterial (Campbell et al. 2013) and archaeal (Rinke et al. 2013) single-copy core genes using HMMER v3.2.1 (Eddy 2011), (3) ‘anvi-run-ncbi-cogs’ to annotate ORFs with functions from the NCBI’s Clusters of Orthologous Groups (COGs) (Tatusov et al. 2003), and (4) ‘anvi-run-kegg-kofams’ to annotate ORFs with functions from the KOfam HMM database of KEGG orthologs (KOs) (Aramaki et al. 2020; M. Kanehisa and Goto 2000). To predict the approximate number of genomes in metagenomic assemblies we used the program ‘anvi-display-contigs-stats’, which calculates the mode of the frequency of single-copy core genes as described previously (Delmont and Eren 2016).

### Metagenomic read recruitment, reconstructing genomes from metagenomes, determination of genome taxonomy and ANI

We recruited metagenomic short reads to contigs using Bowtie2 v2.3.5 (Langmead and Salzberg 2012) and converted resulting SAM files to BAM files using samtools v1.9 (Li et al. 2009). We profiled the resulting BAM files using the program ‘anvi-profile’ with the flag ‘--min-contig-length’ set to 2500 to eliminate shorter sequences to minimize noise. We then used the program ‘anvi-merge’ to combine all read recruitment profiles into a single anvi’o merged profile database for downstream visualization, binning, and statistical analyses (the DOI 10.6084/m9.figshare.14331236 gives access to reproducible data objects). We then used ‘anvi-cluster-contigs’ to group contigs into 100 initial bins using CONCOCT v1.1.0 (Alneberg et al. 2014), ‘anvi-refine’ to manually curate initial bins with conflation error based on tetranucleotide frequency and differential coverage signal across all samples, and ‘anvi-summarize’ to report final summary statistics for each gene, contig, and bin.

We used the program ‘anvi-rename-bins’ to identify bins that were more than 70% complete and less than 10% redundant and store them in a new collection as metagenome-assembled genomes (MAGs), discarding lower quality bins from downstream analyses. GTBD-tk v0.3.2 (Chaumeil et al. 2019) assigned taxonomy to each of our MAGs using GTDB r89 (Parks et al. 2018), but to assign species- and subspecies-level taxonomy for ‘DA_MAG_00057’, ‘DA_MAG_00011’, ‘DA_MAG_00052’ and ‘DA_MAG_00018’, we used ‘anvi-get-sequences-for-hmm-hits’ to recover DNA sequences for bacterial single-copy core genes that encode ribosomal proteins, and searched them in the NCBI’s nucleotide collection (nt) database using BLAST (Altschul et al. 1990). Finally, the program ‘anvi-compute-genome-similarity’ calculated pairwise genomic average nucleotide identity (gANI) of our genomes using PyANI v0.2.9 (Pritchard et al. 2016).

### Criteria for MAG detection in metagenomes

Using mean coverage to assess the occurrence of populations in a given sample based on metagenomic read recruitment can yield misleading insights, since this strategy cannot accurately distinguish reference sequences that represent very low-abundance environmental populations from those sequences that do not represent an environmental population in a sample yet still recruit reads from non-target populations due to the presence of conserved genomic regions. Thus, we relied upon the ‘detection’ metric, which is a measure of the proportion of the nucleotides in a given sequence that are covered by at least one short read. We considered a population to be detected in a metagenome if anvi’o reported a detection value of at least 0.25 for its genome (whether it was a metagenome-assembled or isolate genome). Values of detection in metagenomic read recruitment results often follow a bimodal distribution for populations that are present and absent (see Supplementary Figure 2 in ref. (Utter et al. 2020)), thus 0.25 is an appropriate cutoff to eliminate false-positive signal in read recruitment results for populations that are absent.

### Identification of MAGs that represent multiple subpopulations

To identify subpopulations of MAGs in metagenomes, we used the anvi’o command ‘anvi-gen-variability-profile’ with the ‘--quince-mode’ flag which exported single-nucleotide variant (SNV) information for all MAGs after read recruitment. We then used DESMAN v2.1.1 (Quince et al. 2017) to analyze SNVs to determine the number and distribution of subpopulations represented by a single genome. To account for non-specific mapping that can inflate the number of estimated subpopulations, we removed any subpopulation that made up less than 1% of the entire population explained by a single MAG. To account for noise due to low coverage, we only investigated subpopulations for MAGs for which the mean non-outlier coverage of single-copy core genes was at least 10X.

### Criteria for colonization of a recipient by a MAG for colonization dynamics analyses (Supplementary Information)

We applied the set of criteria described in Supplementary Figure 4 to determine whether or not a MAG successfully colonized a recipient, and to confidently assign colonization or non-colonization phenotypes to each MAG/recipient pair where the MAG was detected in the donor sample used for transplant into the recipient. If these criteria were met, we then determined whether the MAG was detected in any post-FMT recipient sample taken more than 7 days after transplant. If not, the MAG/recipient pair was considered a non-colonization event. If the MAG was detected in the recipient greater than 7 days post-FMT, we used subpopulation information to determine if any subpopulation present in the donor and absent in the recipient pre-FMT was detected in the recipient more than 7 days post-FMT. If this was the case, we considered this to represent a colonization event. See Supplementary Figure 4 for a complete outline of all possible cases.

### Phylogenomic tree construction

To concatenate and align amino acid sequences of 46 single-copy core (Campbell et al. 2013) ribosomal proteins that were present in all of our *Bifidobacterium* MAGs and reference genomes, we ran the anvi’o command ‘anvi-get-sequences-for-hmm-hits’ with the ‘--return-best-hit’, ‘--get-aa-sequence’ and ‘--concatenate’ flags, and the ‘--align-with’ flag set to ‘muscle’ to use MUSCLE v3.8.1551 (Edgar 2004) for alignment. We then ran ‘anvi-gen-phylogenomic-tree’ with default parameters to compute a phylogenomic tree using FastTree 2.1 (Price, Dehal, and Arkin 2010).

### Analysis of metabolic modules and enrichment

We calculated the level of completeness for a given KEGG module (Minoru Kanehisa et al. 2014, 2017) in our genomes using the program ‘anvi-estimate-metabolism’, which leveraged previous annotation of genes with KEGG orthologs (KOs) (see the section ‘Processing of contigs’). Then, the program ‘anvi-compute-functional-enrichment’ determined whether a given metabolic module was enriched in a group of genomes based on the output from the program ‘anvi-estimate-metabolism’. The URL https://anvio.org/help/7.1/programs/anvi-estimate-metabolism/ serves a tutorial for this program which details the modes of usage and output file formats. The statistical approach for enrichment analysis is defined elsewhere (Shaiber et al. 2020), but briefly it computes enrichment scores for functions (or metabolic modules) within groups by fitting a binomial generalized linear model (GLM) to the occurrence of each function or complete metabolic module in each group, and then computing a Rao test statistic, uncorrected p-values, and corrected q-values. We considered any function or metabolic module with a q-value less than 0.05 to be ‘enriched’ in its associated group if it was also at least 75% complete and present in at least 50% of the group members.

### Determination of MAGs representing good and poor colonizers for metabolic enrichment analysis

We classified MAGs as good colonizers if, in all 5 recipients, they were detected in the donor sample used for transplantation as well as the recipient more than 7 days post-FMT. We classified MAGs as poor colonizers as those that, in at least 3 recipients, were detected in the donor sample used for FMT but were not detected in the recipient at least 7 days post-FMT. We reduced the number of good colonizer MAGs to be the same as the number of poor colonizer MAGs for metabolic enrichment analysis by selecting only those populations that were the most prevalent in the Canadian gut metagenomes.

### Ordination plots

We used the R vegan v2.4-2 package ‘metaMDS’ function to perform nonmetric multidimensional scaling (NMDS) with Horn-Morisita dissimilarity distance to compare taxonomic composition between donor, recipient, and global metagenomes. We visualized ordination plots using R ggplot2.

## Code and Data Availability

Raw sequencing data for donor and recipient metagenomes are stored under the NCBI BioProject PRJNA701961 (see Supplementary Table 1 for accession numbers for each sample). The URL https://merenlab.org/data/fmt-gut-colonization serves a reproducible bioinformatics workflow and gives access to ad hoc scripts, usage instructions, and intermediate data objects to reproduce findings in our study. Supplementary tables are accessible also via doi:10.6084/m9.figshare.14138405.

## Acknowledgements

We thank Mitchell L. Sogin, Eugene B. Chang, Samuel H. Light, and Howard A. Shuman for helpful discussions, Ryan Moore and Ozcan C. Esen for technical support, and Nicola Segata and the members of the Segata group for their assistance with genomes from healthy gut metagenomes. We also thank Kaiyu Wu, Robyn Louie and Linda Ward of the IPC Research Laboratory at the University of Calgary for their help with patient recruitment and sampling. This project was supported by the GI Research Foundation (GIRF) and the Mutchnik Family Fund. Additionally, ARW acknowledges support from the Robert C. and Mary Jane Gallo Scholarship Fund; JF acknowledges support from the Alissa and Gianna Carlino Fellowship in Celiac Disease Research; BJ acknowledges support from the Cancer Center Support grant P30CA014599 and Digestive Diseases Research Core Center P30 DK42086; and IV acknowledges support from the National Science Foundation Graduate Research Fellowship (1746045).

## Author Contributions

AME, TL, BJ conceived the study. JZD, MS, DK, TL recruited patients, performed transplantation experiments, and collected samples. ARW, JF, AME performed primary data analyses. IV developed research tools. KL, STML, HGM performed sample processing and sequencing. FT, AS, EF, JMR, CQ, MKY, AY contributed to data analyses and interpretation. DTR, BJ, TL, and AME directed research. ARW, JF, AME wrote the paper with critical input from all authors.

## Supplementary Figures

**Supplementary Figure 1.**
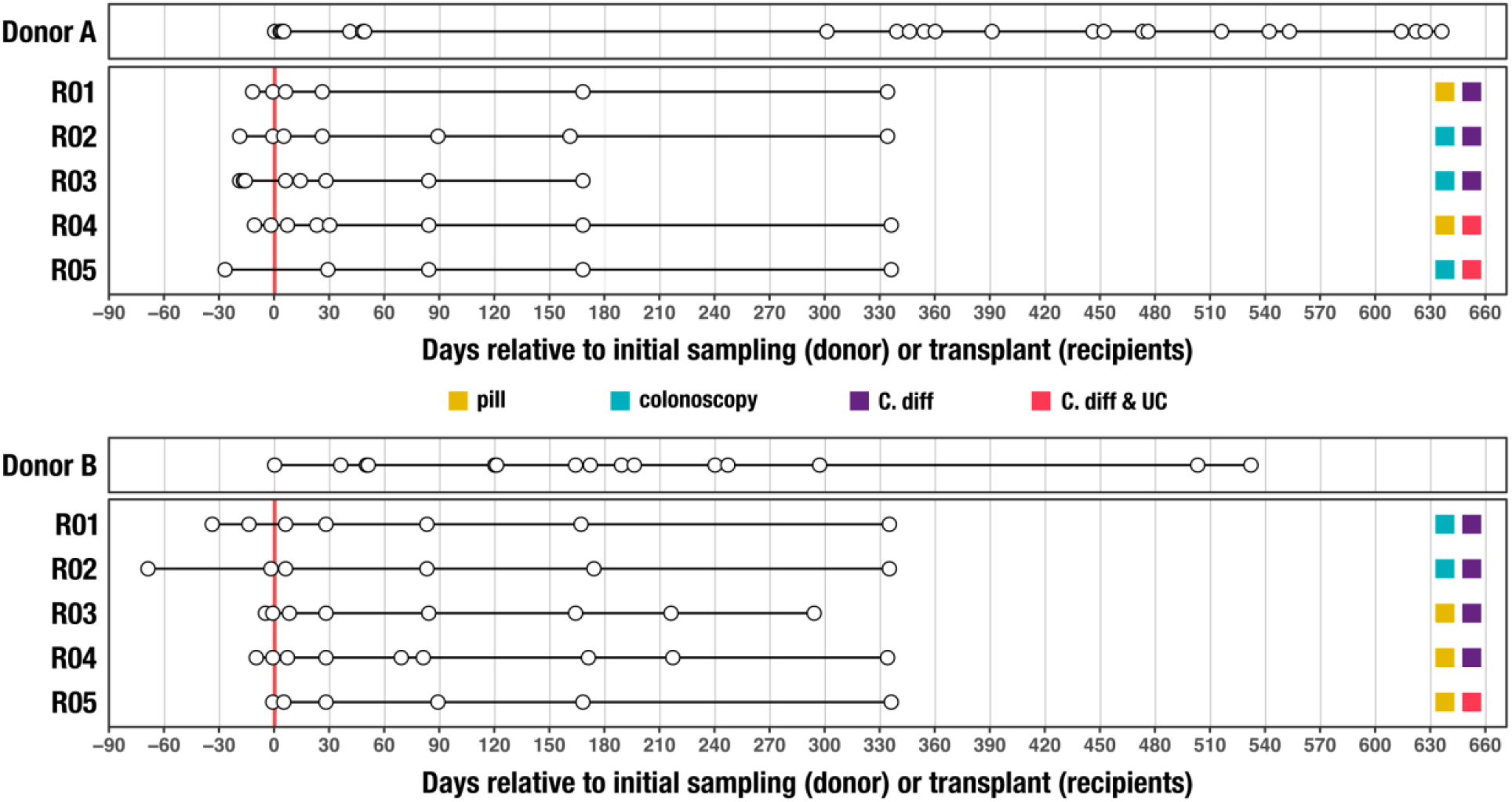
Timeline of stool samples collected from FMT study. Each circle represents a stool sample collected from either an FMT donor or FMT recipient. The thicker, red vertical line at day 0 represents the FMT event for each recipient. FMT method (pill or colonoscopy) and FMT recipient health and disease state (C. diff - chronic recurrent *Clostridium difficile* infection, UC - ulcerative colitis) are indicated on the right.

**Supplementary Figure 2.**
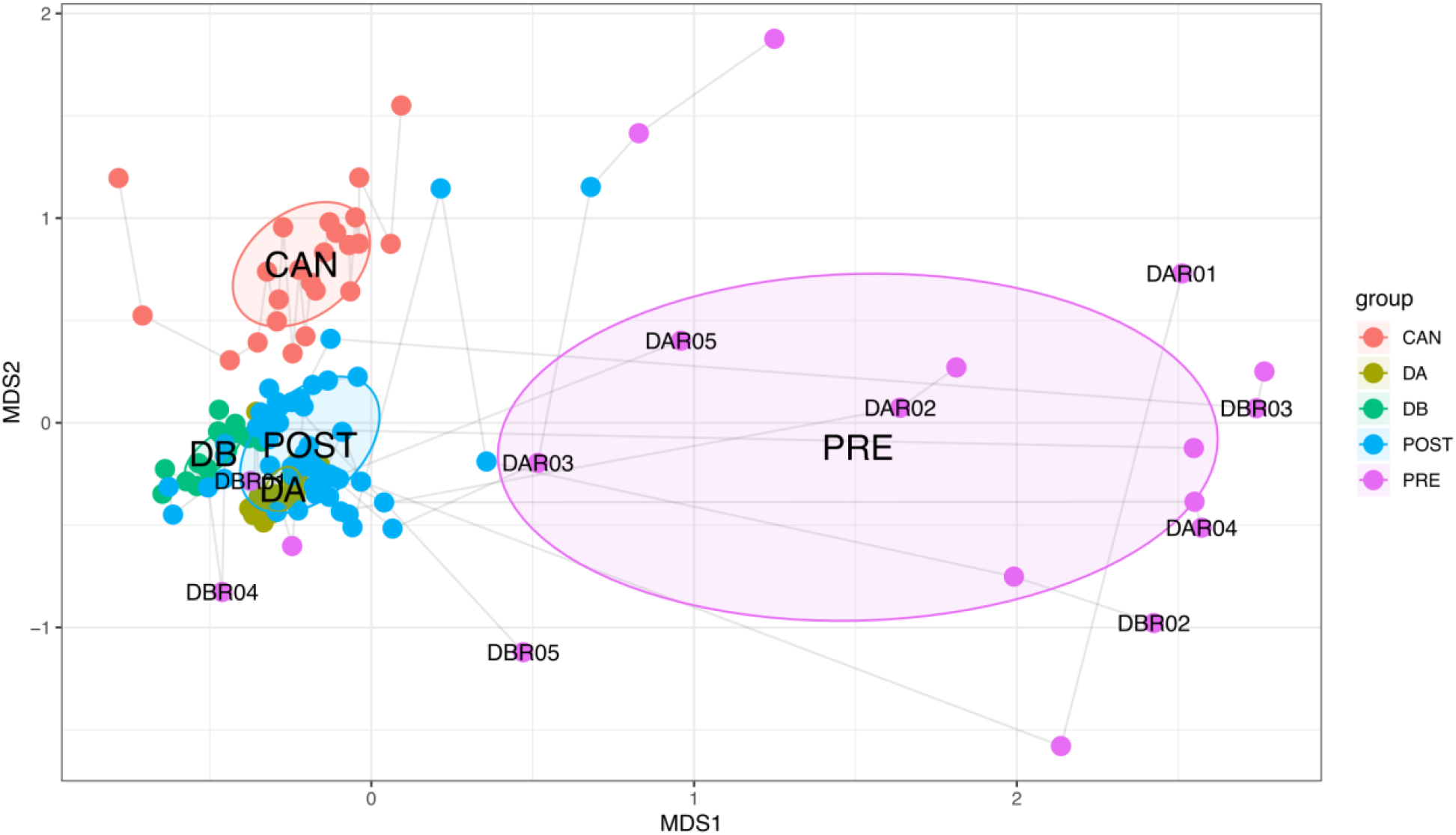
Nonmetric multidimensional scaling (NMDS) ordination of the taxonomic composition of donor, recipient, and Canadian gut metagenomes at the genus level based on Morisita-Horn dissimilarity. Samples from the same participant are joined by lines with the earliest time point labeled. CAN: Canadian gut metagenomes, DA: donor A, DB: donor B, POST: recipients post-FMT, PRE: recipients pre-FMT.

**Supplementary Figure 3.**
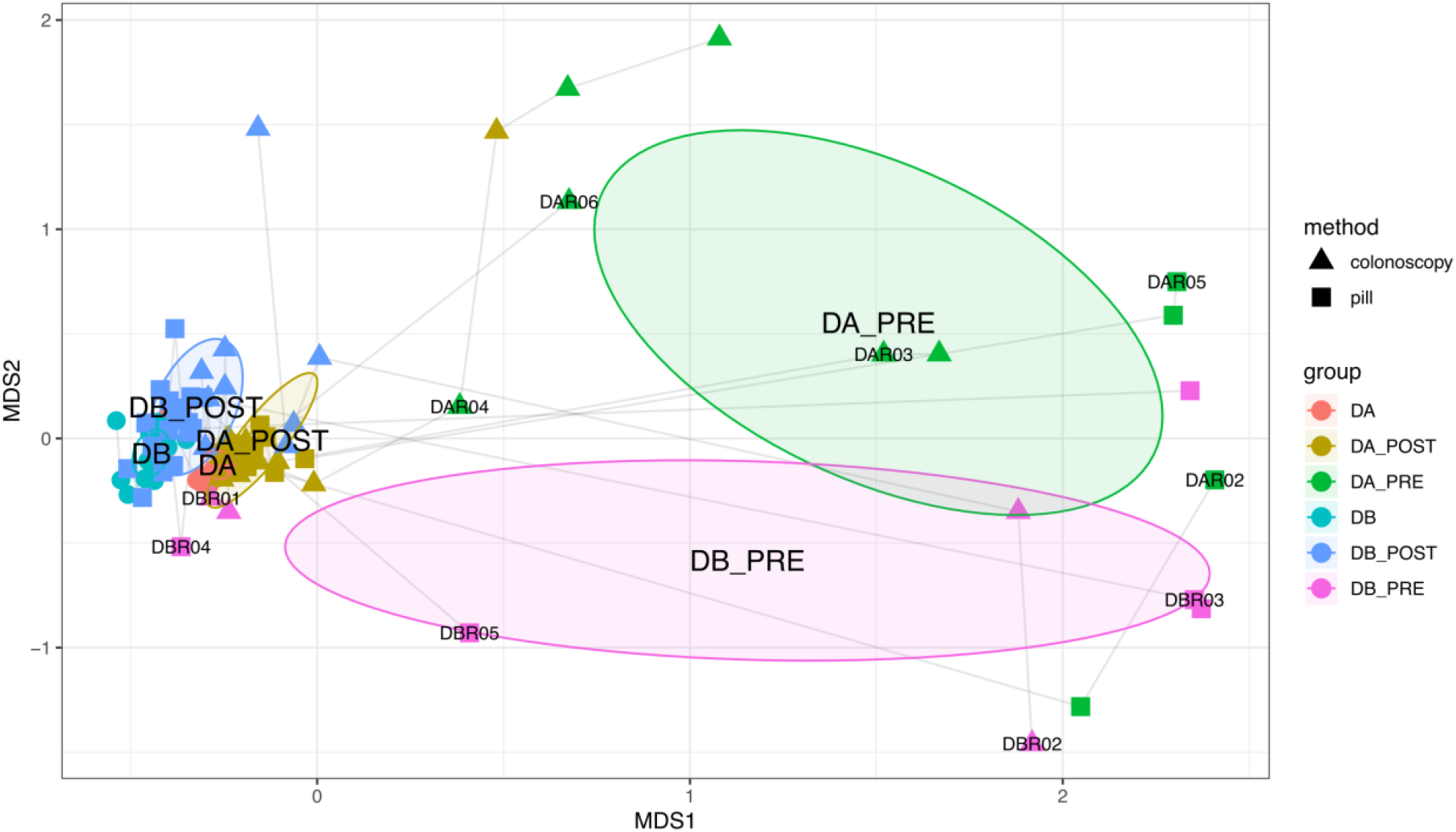
Nonmetric multidimensional scaling (NMDS) ordination of the taxonomic composition of the donor and recipient metagenomes at genus level based on Morisita-Horn dissimilarity. Samples from the same participant are joined by lines with the earliest time point labeled. DA_POST: donor A recipients post-FMT, DA_PRE: donor A recipients pre-FMT, DA: donor A, DB_POST: donor B recipients post-FMT, DB_PRE: donor B recipients pre-FMT, DB: donor B.

**Supplementary Figure 4.**
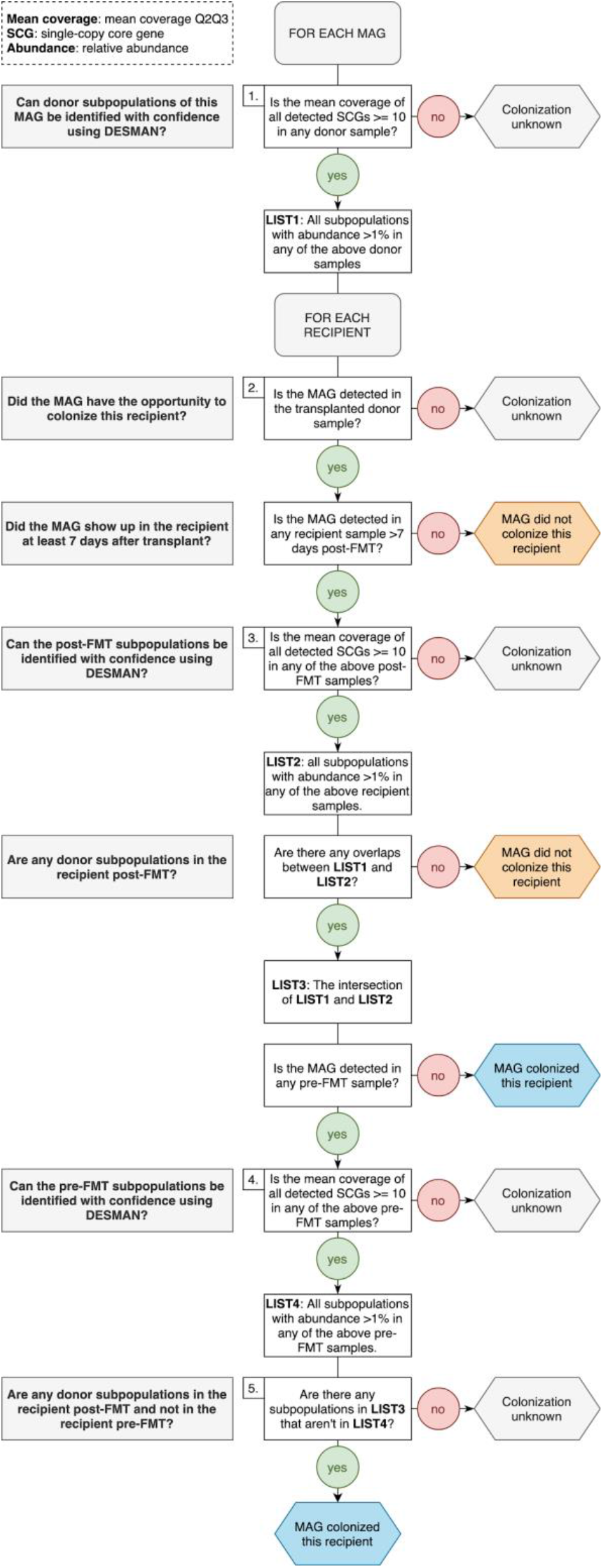
A flowchart outlining our method to assign successful colonization, failed colonization, or undetermined colonization phenotypes to donor-derived populations in the recipients of that donor’s stool.

## Supplementary Tables 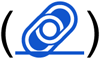

**Supplementary Table 1: Description of FMT study and stool samples collected**. a) Description of FMT donor stool samples and SRA accession numbers. b) Description of FMT recipient samples and SRA accession numbers. c) Description of transplantation events.

**Supplementary Table 2: Description of FMT metagenomes and co-assemblies**. a) Metagenome SRA accession numbers and numbers of metagenomic short-reads sequenced and mapped to co-assemblies and MAGs. b) Phylum level taxonomic composition of metagenomes. c) Genus level taxonomic composition of metagenomes. d) Summary statistics for contigs from metagenome co-assemblies.

**Supplementary Table 3: Description of MAGs**. a) Summary statistics and taxonomic assignments for MAGs. b) and c) Detection of Donor A and Donor B MAGs in FMT metagenomes, respectively. d) and e) Detection of Donor A and Donor B MAGs in global gut metagenomes, respectively. f) and g) Detection summary statistics of Donor A and Donor B MAGs in global gut metagenomes, respectively. h) and i) Mean non-outlier coverage of Donor A and Donor B MAG single-copy core genes in FMT metagenomes.

**Supplementary Table 4:** Accession numbers of gut metagenomes from 17 countries.

**Supplementary Table 5: MAG subpopulation information**. a) and b) Number of Donor A and Donor B MAG subpopulations detected in FMT metagenomes, respectively. c) and d) Subpopulation composition of Donor A and Donor B MAGs in FMT metagenomes, respectively.

**Supplementary Table 6:** MAG/recipient pair colonization outcomes and MAG mean coverage in the 2nd and 3rd quartiles in stool samples used for transplantation.

**Supplementary Table 7: Description of HMI vs. LMI populations**. a) Taxonomic assignments and genome size estimates for high- and low-metabolic independence populations. b) KEGG module completeness information for high- and low-metabolic independence populations. c) Raw KEGG module enrichment information for high- and low-metabolic independence populations. d) KEGG module enrichment and categorical information for the 33 modules enriched in high-metabolic independence populations. e) and f) Completeness information for the 33 modules enriched in high-metabolic independence populations in all high- and low-metabolic independence populations.

**Supplementary Table 8:** a) List of genomes from healthy individuals and individuals with IBD. b) Module completion values across genomes.

## References

Almeida, Cátia, Rita Oliveira, Raquel Soares, and Pedro Barata. 2020. “Influence of Gut Microbiota Dysbiosis on Brain Function: A Systematic Review.” Porto Biomedical Journal 5 (2). https://doi.org/10.1097/j.pbj.0000000000000059.

Alneberg, Johannes, Brynjar Smári Bjarnason, Ino de Bruijn, Melanie Schirmer, Joshua Quick, Umer Z. Ijaz, Leo Lahti, Nicholas J. Loman, Anders F. Andersson, and Christopher Quince. 2014. “Binning Metagenomic Contigs by Coverage and Composition.” Nature Methods 11 (11): 1144–46.

Altschul, S. F., W. Gish, W. Miller, E. W. Myers, and D. J. Lipman. 1990. “Basic Local Alignment Search Tool.” Journal of Molecular Biology 215 (3): 403–10.

Aramaki, Takuya, Romain Blanc-Mathieu, Hisashi Endo, Koichi Ohkubo, Minoru Kanehisa, Susumu Goto, and Hiroyuki Ogata. 2020. “KofamKOALA: KEGG Ortholog Assignment Based on Profile HMM and Adaptive Score Threshold.” Bioinformatics 36 (7): 2251–52.

Arumugam, Manimozhiyan, Jeroen Raes, Eric Pelletier, Denis Le Paslier, Takuji Yamada, Daniel R. Mende, Gabriel R. Fernandes, et al. 2011. “Enterotypes of the Human Gut Microbiome.” Nature 473 (7346): 174–80.

Bäckhed, Fredrik, Claire M. Fraser, Yehuda Ringel, Mary Ellen Sanders, R. Balfour Sartor, Philip M. Sherman, James Versalovic, Vincent Young, and B. Brett Finlay. 2012. “Defining a Healthy Human Gut Microbiome: Current Concepts, Future Directions, and Clinical Applications.” Cell Host & Microbe 12 (5): 611–22.

Baumgart, Daniel C., and Simon R. Carding. 2007. “Inflammatory Bowel Disease: Cause and Immunobiology.” The Lancet 369 (9573): 1627–40.

Biesalski, Hans K. 2016. “Nutrition Meets the Microbiome: Micronutrients and the Microbiota.” Annals of the New York Academy of Sciences 1372 (1): 53–64.

Bowers, Robert M., Nikos C. Kyrpides, Ramunas Stepanauskas, Miranda Harmon-Smith, Devin Doud, T. B. K. Reddy, Frederik Schulz, et al. 2017. “Minimum Information about a Single Amplified Genome (MISAG) and a Metagenome-Assembled Genome (MIMAG) of Bacteria and Archaea.” Nature Biotechnology 35 (8): 725–31.

Campbell, James H., Patrick O’Donoghue, Alisha G. Campbell, Patrick Schwientek, Alexander Sczyrba, Tanja Woyke, Dieter Söll, and Mircea Podar. 2013. “UGA Is an Additional Glycine Codon in Uncultured SR1 Bacteria from the Human Microbiota.” Proceedings of the National Academy of Sciences of the United States of America 110 (14): 5540–45.

Chaumeil, Pierre-Alain, Aaron J. Mussig, Philip Hugenholtz, and Donovan H. Parks. 2019. “GTDB-Tk: A Toolkit to Classify Genomes with the Genome Taxonomy Database.” Bioinformatics, November. https://doi.org/10.1093/bioinformatics/btz848.

Chen, Lin-Xing, Karthik Anantharaman, Alon Shaiber, A. Murat Eren, and Jillian F. Banfield. 2020. “Accurate and Complete Genomes from Metagenomes.” Genome Research 30 (3): 315–33.

Chow, Janet, Haiqing Tang, and Sarkis K. Mazmanian. 2011. “Pathobionts of the Gastrointestinal Microbiota and Inflammatory Disease.” Current Opinion in Immunology 23 (4): 473–80.

Clooney, Adam G., Julia Eckenberger, Emilio Laserna-Mendieta, Kathryn A. Sexton, Matthew T. Bernstein, Kathy Vagianos, Michael Sargent, et al. 2021. “Ranking Microbiome Variance in Inflammatory Bowel Disease: A Large Longitudinal Intercontinental Study.” Gut 70 (3): 499–510.

Costello, Elizabeth K., Keaton Stagaman, Les Dethlefsen, Brendan J. M. Bohannan, and David A. Relman. 2012. “The Application of Ecological Theory toward an Understanding of the Human Microbiome.” Science 336 (6086): 1255–62.

David, Lawrence A., Arne C. Materna, Jonathan Friedman, Maria I. Campos-Baptista, Matthew C. Blackburn, Allison Perrotta, Susan E. Erdman, and Eric J. Alm. 2014. “Host Lifestyle Affects Human Microbiota on Daily Timescales.” Genome Biology 15 (7): R89.

Delmont, Tom O., and A. Murat Eren. 2016. “Identifying Contamination with Advanced Visualization and Analysis Practices: Metagenomic Approaches for Eukaryotic Genome Assemblies.” PeerJ 4 (March): e1839.

Delmont, Tom O., Christopher Quince, Alon Shaiber, Özcan C. Esen, Sonny Tm Lee, Michael S. Rappé, Sandra L. McLellan, Sebastian Lücker, and A. Murat Eren. 2018. “Nitrogen-Fixing Populations of Planctomycetes and Proteobacteria Are Abundant in Surface Ocean Metagenomes.” Nature Microbiology 3 (7): 804–13.

De Preter, Vicky, Veerle Bulteel, Peter Suenaert, Karen Paula Geboes, Gert De Hertogh, Anja Luypaerts, Karel Geboes, Kristin Verbeke, and Paul Rutgeerts. 2009. “Pouchitis, Similar to Active Ulcerative Colitis, Is Associated with Impaired Butyrate Oxidation by Intestinal Mucosa.” Inflammatory Bowel Diseases 15 (3): 335–40.

Donaldson, Gregory P., S. Melanie Lee, and Sarkis K. Mazmanian. 2016. “Gut Biogeography of the Bacterial Microbiota.” Nature Reviews. Microbiology 14 (1): 20–32.

D’Souza, Glen, Shraddha Shitut, Daniel Preussger, Ghada Yousif, Silvio Waschina, and Christian Kost. 2018. “Ecology and Evolution of Metabolic Cross-Feeding Interactions in Bacteria.” Natural Product Reports 35 (5): 455–88.

Durack, Juliana, and Susan V. Lynch. 2019. “The Gut Microbiome: Relationships with Disease and Opportunities for Therapy.” The Journal of Experimental Medicine 216 (1): 20–40.

Eddy, Sean R. 2011. “Accelerated Profile HMM Searches.” PLoS Computational Biology 7 (10): e1002195.

Edgar, Robert C. 2004. “MUSCLE: Multiple Sequence Alignment with High Accuracy and High Throughput.” Nucleic Acids Research 32 (5): 1792–97.

Eiseman, B., W. Silen, G. S. Bascom, and A. J. Kauvar. 1958. “Fecal Enema as an Adjunct in the Treatment of Pseudomembranous Enterocolitis.” Surgery 44 (5): 854–59.

Eisenstein, Michael. 2020. “The Hunt for a Healthy Microbiome.” Nature 577 (7792): S6–8.

Eren, A. Murat, Özcan C. Esen, Christopher Quince, Joseph H. Vineis, Hilary G. Morrison, Mitchell L. Sogin, and Tom O. Delmont. 2015. “Anvi’o: An Advanced Analysis and Visualization Platform for ‘Omics Data.” PeerJ 3 (October): e1319.

Eren, A. Murat, Evan Kiefl, Alon Shaiber, Iva Veseli, Samuel E. Miller, Matthew S. Schechter, Isaac Fink, et al. 2021. “Community-Led, Integrated, Reproducible Multi-Omics with Anvi’o.” Nature Microbiology 6 (1): 3–6.

Eren, A. Murat, Joseph H. Vineis, Hilary G. Morrison, and Mitchell L. Sogin. 2013. “A Filtering Method to Generate High Quality Short Reads Using Illumina Paired-End Technology.” PloS One 8 (6): e66643.

Feng, Lihui, Arjun S. Raman, Matthew C. Hibberd, Jiye Cheng, Nicholas W. Griffin, Yangqing Peng, Semen A. Leyn, Dmitry A. Rodionov, Andrei L. Osterman, and Jeffrey I. Gordon. 2020. “Identifying Determinants of Bacterial Fitness in a Model of Human Gut Microbial Succession.” Proceedings of the National Academy of Sciences of the United States of America 117 (5): 2622–33.

Grehan, Martin J., Thomas Julius Borody, Sharyn M. Leis, Jordana Campbell, Hazel Mitchell, and Antony Wettstein. 2010. “Durable Alteration of the Colonic Microbiota by the Administration of Donor Fecal Flora.” Journal of Clinical Gastroenterology 44 (8): 551–61.

Herrmann, Klaus M., and Lisa M. Weaver. 1999. “THE SHIKIMATE PATHWAY.” Annual Review of Plant Physiology and Plant Molecular Biology 50 (June): 473–503.

Human Microbiome Project Consortium. 2012. “Structure, Function and Diversity of the Healthy Human Microbiome.” Nature 486 (7402): 207–14.

Hyatt, Doug, Gwo-Liang Chen, Philip F. Locascio, Miriam L. Land, Frank W. Larimer, and Loren J. Hauser. 2010. “Prodigal: Prokaryotic Gene Recognition and Translation Initiation Site Identification.” BMC Bioinformatics 11 (March): 119.

Isaac, Sandrine, Jose U. Scher, Ana Djukovic, Nuria Jiménez, Dan R. Littman, Steven B. Abramson, Eric G. Pamer, and Carles Ubeda. 2017. “Short- and Long-Term Effects of Oral Vancomycin on the Human Intestinal Microbiota.” The Journal of Antimicrobial Chemotherapy 72 (1): 128–36.

Jimenez, Miguel, Robert Langer, and Giovanni Traverso. 2019. “Microbial Therapeutics: New Opportunities for Drug Delivery.” The Journal of Experimental Medicine 216 (5): 1005–9.

Joossens, Marie, Geert Huys, Margo Cnockaert, Vicky De Preter, Kristin Verbeke, Paul Rutgeerts, Peter Vandamme, and Severine Vermeire. 2011. “Dysbiosis of the Faecal Microbiota in Patients with Crohn’s Disease and Their Unaffected Relatives.” Gut 60 (5): 631–37.

Kanehisa, M., and S. Goto. 2000. “KEGG: Kyoto Encyclopedia of Genes and Genomes.” Nucleic Acids Research 28 (1): 27–30.

Kanehisa, Minoru, Miho Furumichi, Mao Tanabe, Yoko Sato, and Kanae Morishima. 2017. “KEGG: New Perspectives on Genomes, Pathways, Diseases and Drugs.” Nucleic Acids Research 45 (D1): D353–61.

Kanehisa, Minoru, Susumu Goto, Yoko Sato, Masayuki Kawashima, Miho Furumichi, and Mao Tanabe. 2014. “Data, Information, Knowledge and Principle: Back to Metabolism in KEGG.” Nucleic Acids Research 42 (Database issue): D199–205.

Kao, Dina, Brandi Roach, Marisela Silva, Paul Beck, Kevin Rioux, Gilaad G. Kaplan, Hsiu-Ju Chang, et al. 2017. “Effect of Oral Capsule-vs Colonoscopy-Delivered Fecal Microbiota Transplantation on Recurrent Clostridium Difficile Infection: A Randomized Clinical Trial.” JAMA: The Journal of the American Medical Association 318 (20): 1985–93.

Khoruts, Alexander, Johan Dicksved, Janet K. Jansson, and Michael J. Sadowsky. 2010. “Changes in the Composition of the Human Fecal Microbiome after Bacteriotherapy for Recurrent Clostridium Difficile-Associated Diarrhea.” Journal of Clinical Gastroenterology 44 (5): 354–60.

Koenig, Jeremy E., Aymé Spor, Nicholas Scalfone, Ashwana D. Fricker, Jesse Stombaugh, Rob Knight, Largus T. Angenent, and Ruth E. Ley. 2011. “Succession of Microbial Consortia in the Developing Infant Gut Microbiome.” Proceedings of the National Academy of Sciences of the United States of America 108 Suppl 1 (March): 4578–85.

Koropatkin, Nicole M., Elizabeth A. Cameron, and Eric C. Martens. 2012. “How Glycan Metabolism Shapes the Human Gut Microbiota.” Nature Reviews. Microbiology 10 (5): 323–35.

Köster, Johannes, and Sven Rahmann. 2012. “Snakemake—a Scalable Bioinformatics Workflow Engine.” Bioinformatics 28 (19): 2520–22.

Langmead, Ben, and Steven L. Salzberg. 2012. “Fast Gapped-Read Alignment with Bowtie 2.” Nature Methods 9 (4): 357–59.

Lee, Sonny T. M., Stacy A. Kahn, Tom O. Delmont, Alon Shaiber, Özcan C. Esen, Nathaniel A. Hubert, Hilary G. Morrison, Dionysios A. Antonopoulos, David T. Rubin, and A. Murat Eren. 2017. “Tracking Microbial Colonization in Fecal Microbiota Transplantation Experiments via Genome-Resolved Metagenomics.” Microbiome 5 (1): 50.

Li, Heng, Bob Handsaker, Alec Wysoker, Tim Fennell, Jue Ruan, Nils Homer, Gabor Marth, Goncalo Abecasis, Richard Durbin, and 1000 Genome Project Data Processing Subgroup. 2009. “The Sequence Alignment/Map Format and SAMtools.” Bioinformatics 25 (16): 2078–79.

Lloyd-Price, Jason, Galeb Abu-Ali, and Curtis Huttenhower. 2016. “The Healthy Human Microbiome.” Genome Medicine 8 (1): 51.

Lloyd-Price, Jason, Cesar Arze, Ashwin N. Ananthakrishnan, Melanie Schirmer, Julian Avila-Pacheco, Tiffany W. Poon, Elizabeth Andrews, et al. 2019. “Multi-Omics of the Gut Microbial Ecosystem in Inflammatory Bowel Diseases.” Nature 569 (7758): 655–62.

Lynch, Susan V., and Oluf Pedersen. 2016. “The Human Intestinal Microbiome in Health and Disease.” The New England Journal of Medicine 375 (24): 2369–79.

Martens, Eric C., Herbert C. Chiang, and Jeffrey I. Gordon. 2008. “Mucosal Glycan Foraging Enhances Fitness and Transmission of a Saccharolytic Human Gut Bacterial Symbiont.” Cell Host & Microbe 4 (5): 447–57.

Martens, J. H., H. Barg, M. J. Warren, and D. Jahn. 2002. “Microbial Production of Vitamin B12.” Applied Microbiology and Biotechnology 58 (3): 275–85.

McBurney, Michael I., Cindy Davis, Claire M. Fraser, Barbara O. Schneeman, Curtis Huttenhower, Kristin Verbeke, Jens Walter, and Marie E. Latulippe. 2019. “Establishing What Constitutes a Healthy Human Gut Microbiome: State of the Science, Regulatory Considerations, and Future Directions.” The Journal of Nutrition 149 (11): 1882–95.

Messer, J. S., E. R. Liechty, O. A. Vogel, and E. B. Chang. 2017. “Evolutionary and Ecological Forces That Shape the Bacterial Communities of the Human Gut.” Mucosal Immunology 10 (3): 567–79.

Minoche, André E., Juliane C. Dohm, and Heinz Himmelbauer. 2011. “Evaluation of Genomic High-Throughput Sequencing Data Generated on Illumina HiSeq and Genome Analyzer Systems.” Genome Biology 12 (11): R112.

Nood, Els van, Anne Vrieze, Max Nieuwdorp, Susana Fuentes, Erwin G. Zoetendal, Willem M. de Vos, Caroline E. Visser, et al. 2013. “Duodenal Infusion of Donor Feces for Recurrent Clostridium Difficile.” The New England Journal of Medicine 368 (5): 407–15.

Ott, S. J., M. Musfeldt, D. F. Wenderoth, J. Hampe, O. Brant, U. R. Fölsch, K. N. Timmis, and S. Schreiber. 2004. “Reduction in Diversity of the Colonic Mucosa Associated Bacterial Microflora in Patients with Active Inflammatory Bowel Disease.” Gut 53 (5): 685–93.

Parks, Donovan H., Maria Chuvochina, David W. Waite, Christian Rinke, Adam Skarshewski, Pierre-Alain Chaumeil, and Philip Hugenholtz. 2018. “A Standardized Bacterial Taxonomy Based on Genome Phylogeny Substantially Revises the Tree of Life.” Nature Biotechnology 36 (10): 996–1004.

Pasolli, Edoardo, Francesco Asnicar, Serena Manara, Moreno Zolfo, Nicolai Karcher, Federica Armanini, Francesco Beghini, et al. 2019. “Extensive Unexplored Human Microbiome Diversity Revealed by Over 150,000 Genomes from Metagenomes Spanning Age, Geography, and Lifestyle.” Cell 176 (3): 649–62.e20.

Peng, Yu, Henry C. M. Leung, S. M. Yiu, and Francis Y. L. Chin. 2012. “IDBA-UD: A de Novo Assembler for Single-Cell and Metagenomic Sequencing Data with Highly Uneven Depth.” Bioinformatics 28 (11): 1420–28.

Plichta, Damian R., Daniel B. Graham, Sathish Subramanian, and Ramnik J. Xavier. 2019. “Therapeutic Opportunities in Inflammatory Bowel Disease: Mechanistic Dissection of Host-Microbiome Relationships.” Cell 178 (5): 1041–56.

Price, Morgan N., Paramvir S. Dehal, and Adam P. Arkin. 2010. “FastTree 2--Approximately Maximum-Likelihood Trees for Large Alignments.” PloS One 5 (3): e9490.

Pritchard, Leighton, Rachel H. Glover, Sonia Humphris, John G. Elphinstone, and Ian K. Toth. 2016. “Genomics and Taxonomy in Diagnostics for Food Security: Soft-Rotting Enterobacterial Plant Pathogens.” Analytical Methods 8 (1): 12–24.

Quince, Christopher, Tom O. Delmont, Sébastien Raguideau, Johannes Alneberg, Aaron E. Darling, Gavin Collins, and A. Murat Eren. 2017. “DESMAN: A New Tool for de Novo Extraction of Strains from Metagenomes.” Genome Biology 18 (1): 181.

Quince, Christopher, Umer Zeeshan Ijaz, Nick Loman, A. Murat Eren, Delphine Saulnier, Julie Russell, Sarah J. Haig, et al. 2015. “Extensive Modulation of the Fecal Metagenome in Children with Crohn’s Disease During Exclusive Enteral Nutrition.” The American Journal of Gastroenterology 110 (12): 1718–29; quiz 1730.

Rinke, Christian, Patrick Schwientek, Alexander Sczyrba, Natalia N. Ivanova, Iain J. Anderson, Jan-Fang Cheng, Aaron Darling, et al. 2013. “Insights into the Phylogeny and Coding Potential of Microbial Dark Matter.” Nature 499 (7459): 431–37.

Rothschild, Daphna, Omer Weissbrod, Elad Barkan, Alexander Kurilshikov, Tal Korem, David Zeevi, Paul I. Costea, et al. 2018. “Environment Dominates over Host Genetics in Shaping Human Gut Microbiota.” Nature 555 (7695): 210–15.

Schirmer, Melanie, Ashley Garner, Hera Vlamakis, and Ramnik J. Xavier. 2019. “Microbial Genes and Pathways in Inflammatory Bowel Disease.” Nature Reviews. Microbiology 17 (8): 497–511.

Schmidt, Thomas S. B., Jeroen Raes, and Peer Bork. 2018. “The Human Gut Microbiome: From Association to Modulation.” Cell 172 (6): 1198–1215.

Shahinas, Dea, Michael Silverman, Taylor Sittler, Charles Chiu, Peter Kim, Emma Allen-Vercoe, Scott Weese, Andrew Wong, Donald E. Low, and Dylan R. Pillai. 2012. “Toward an Understanding of Changes in Diversity Associated with Fecal Microbiome Transplantation Based on 16S rRNA Gene Deep Sequencing.” mBio 3 (5): e00338–12.

Shaiber, Alon, Amy D. Willis, Tom O. Delmont, Simon Roux, Lin-Xing Chen, Abigail C. Schmid, Mahmoud Yousef, et al. 2020. “Functional and Genetic Markers of Niche Partitioning among Enigmatic Members of the Human Oral Microbiome.” Genome Biology 21 (1): 292.

Sharon, Itai, Michael J. Morowitz, Brian C. Thomas, Elizabeth K. Costello, David A. Relman, and Jillian F. Banfield. 2013. “Time Series Community Genomics Analysis Reveals Rapid Shifts in Bacterial Species, Strains, and Phage during Infant Gut Colonization.” Genome Research 23 (1): 111–20.

Sheth, Ravi U., Mingqiang Li, Weiqian Jiang, Peter A. Sims, Kam W. Leong, and Harris H. Wang. 2019. “Spatial Metagenomic Characterization of Microbial Biogeography in the Gut.” Nature Biotechnology 37 (8): 877–83.

Smillie, Christopher S., Jenny Sauk, Dirk Gevers, Jonathan Friedman, Jaeyun Sung, Ilan Youngster, Elizabeth L. Hohmann, et al. 2018. “Strain Tracking Reveals the Determinants of Bacterial Engraftment in the Human Gut Following Fecal Microbiota Transplantation.” Cell Host & Microbe 23 (2): 229–40.e5.

Sokol, Harry, and Philippe Seksik. 2010. “The Intestinal Microbiota in Inflammatory Bowel Diseases: Time to Connect with the Host.” Current Opinion in Gastroenterology 26 (4): 327–31.

Stewart, Christopher J., Nadim J. Ajami, Jacqueline L. O’Brien, Diane S. Hutchinson, Daniel P. Smith, Matthew C. Wong, Matthew C. Ross, et al. 2018. “Temporal Development of the Gut Microbiome in Early Childhood from the TEDDY Study.” Nature 562 (7728): 583–88.

Swidsinski, Alexander, Jutta Weber, Vera Loening-Baucke, Laura P. Hale, and Herbert Lochs. 2005. “Spatial Organization and Composition of the Mucosal Flora in Patients with Inflammatory Bowel Disease.” Journal of Clinical Microbiology 43 (7): 3380–89.

Tatusov, Roman L., Natalie D. Fedorova, John D. Jackson, Aviva R. Jacobs, Boris Kiryutin, Eugene V. Koonin, Dmitri M. Krylov, et al. 2003. “The COG Database: An Updated Version Includes Eukaryotes.” BMC Bioinformatics 4 (September): 41.

Utter, Daniel R., Gary G. Borisy, A. Murat Eren, Colleen M. Cavanaugh, and Jessica L. Mark Welch. 2020. “Metapangenomics of the Oral Microbiome Provides Insights into Habitat Adaptation and Cultivar Diversity.” Genome Biology 21 (1): 293.

Vanni, Chiara, Matthew S. Schechter, Silvia G. Acinas, Albert Barberán, Pier Luigi Buttigieg, Emilio O. Casamayor, Tom O. Delmont, et al. 2020. “Unifying the Global Coding Sequence Space Enables the Study of Genes with Unknown Function across Biomes.” Cold Spring Harbor Laboratory. https://doi.org/10.1101/2020.06.30.180448.

Vineis, Joseph H., Daina L. Ringus, Hilary G. Morrison, Tom O. Delmont, Sushila Dalal, Laura H. Raffals, Dionysios A. Antonopoulos, et al. 2016. “Patient-Specific Bacteroides Genome Variants in Pouchitis.” mBio 7 (6): e01713.–16, /mbio/7/6/e01713–16.atom.

Walter, Jens, Anissa M. Armet, B. Brett Finlay, and Fergus Shanahan. 2020. “Establishing or Exaggerating Causality for the Gut Microbiome: Lessons from Human Microbiota-Associated Rodents.” Cell 180 (2): 221–32.

Wexler, Aaron G., and Andrew L. Goodman. 2017. “An Insider’s Perspective: Bacteroides as a Window into the Microbiome.” Nature Microbiology. https://doi.org/10.1038/nmicrobiol.2017.26.

Wood, Derrick E., Jennifer Lu, and Ben Langmead. 2019. “Improved Metagenomic Analysis with Kraken 2.” Genome Biology 20 (1): 257.

Wu, Guojun, Naisi Zhao, Chenhong Zhang, Yan Y. Lam, and Liping Zhao. 2021. “Guild-Based Analysis for Understanding Gut Microbiome in Human Health and Diseases.” Genome Medicine 13 (1): 22.

Yasuda, Koji, Keunyoung Oh, Boyu Ren, Timothy L. Tickle, Eric A. Franzosa, Lynn M. Wachtman, Andrew D. Miller, et al. 2015. “Biogeography of the Intestinal Mucosal and Lumenal Microbiome in the Rhesus Macaque.” Cell Host & Microbe 17 (3): 385–91.

Zmora, Niv, Jotham Suez, and Eran Elinav. 2019. “You Are What You Eat: Diet, Health and the Gut Microbiota.” Nature Reviews. Gastroenterology & Hepatology 16 (1): 35–56.

