## Supplementary Information for "Metabolic independence drives gut microbial colonization and resilience in health and disease"

### Supplementary Information for the study “Metabolic independence drives gut microbial colonization and resilience in health and disease”

Andrea R. Watson, Jessika Füssel, Iva Veseli, Johanna Zaal DeLongchamp, Marisela Silva, Florian Trigodet, Karen Lolans, Alon Shaiber, Emily Fogarty, Joseph M. Runde, Christopher Quince, Michael K. Yu, Arda Söylev, Hilary G. Morrison, Sonny T.M. Lee, Dina Kao, David T. Rubin, Bana Jabri, Thomas Louie, A. Murat Eren

*(Please see the main text for author affiliations)*

#### Adaptive ecological forces are the primary drivers of microbial colonization

The ability of a microbial population to colonize and persist in a complex ecosystem is influenced by both neutral and adaptive forces (Maignien et al., 2014). Although which of these is the major driver of successful colonization of the human gut remains unclear (Smillie et al., 2018). In the context of FMT, previous studies have suggested neutral processes to determine colonization success based on the abundance of a microbial population in a donor stool sample (Podlesny and Florian Fricke, 2020; Smillie et al., 2018). Indeed, ecological drift may have a significant role in a system dominated by neutral processes, where low-abundance donor populations in the transplant would be less likely to be observed in recipients. In contrast, if the system is dominated by adaptive forces, colonization success would be a function of the population fitness in the recipient environment, rather than its abundance in the transplant.

To investigate the impact of neutral versus adaptive processes on colonization in our dataset we first asked whether the prevalence of a donor population in publicly available healthy human gut metagenomes, which we define here as a measure of its fitness, was associated with the detection of the same population in donor or recipient metagenomes. Within both FMT cohorts, the mean detection of each population in recipients post-FMT had a stronger association with population fitness than mean detection in donor samples (Figure SIa). The fitness of donor A populations explained 4.2% of the variation in mean detection of those populations in donor samples ($R^2$=0.042, p=0.021) and 19% of variation in mean detection in recipient post-FMT samples ($R^2$=0.19, p=2.7e-07), an increase of approximately 4.5-fold (Figure SIa). Similarly, Donor B population fitness explained 7.3% of the variation in mean detection in donor samples ($R^2$=0.073, p=2.1e-04), and 36% of the variation in mean detection in recipient post-FMT samples ($R^2$=0.36, p=4.5e-19), an increase of approximately 5-fold (Figure SIa). This suggests that fitness is a better predictor of colonization outcome than it is of the detection of a population in the donor, suggesting that adaptive forces are likely at play. But detecting a donor population in a recipient post-FMT metagenome through metagenomic read recruitment does not prove colonization, since donor genomes can recruit reads from recipient populations that are closely related (i.e., strain variants) and that were low abundance prior to FMT.

Resolving colonization events accurately is a challenging task as multiple factors may influence the ability to determine colonization outcomes unambiguously. These factors include (1) the inability to detect low-abundance populations, (2) inaccurate characterization of transient populations observed immediately after FMT as successful colonization events, (3) the reliance on relative abundance of populations to define colonization events when abundance estimates from stool do not always reflect the abundance of organisms in the GI tract (Yasuda et al. 2015; Sheth et al. 2019), and (4) the failure to distinguish between colonization by a donor population or emergence of a pre-FMT recipient population after FMT (where a low-abundance recipient population that is closely related to one or more donor populations becomes abundant after FMT and is mistaken as a bona fide colonization event). To mitigate these factors, we have (1) employed deep-sequencing of our metagenomes which averaged 71 million reads per sample, (2) implemented a longitudinal sampling strategy, that spanned 376 days on average, to observe donor populations in our recipients for an extended period after FMT, (3) leveraged a ‘detection’ metric to define colonization events by presence/absence of populations rather than abundance, and (4) employed microbial population genetics to identify and resolve origins of subpopulations using single-nucleotide variants in read recruitment results (Denef, 2019) and quantify their dynamics (Quince et al., 2017). We also developed an analytical approach that took into consideration the presence and absence of distinct subpopulations in the transplant sample, in the recipient pre-FMT, and in the recipient post-FMT (see Materials and Methods, Supplementary Table 5) to determine whether a given donor population has colonized a given recipient. To determine colonization outcomes, we analyzed 640 genome/recipient pairs for Donor A (128 donor genomes in 5 recipients) and identified 99 successful colonization events, 38 failed colonization events, and 503 ambiguous colonization events (Supplementary Table 6). For Donor B, we analyzed 915 genome/recipient pairs (183 donor genomes in 5 recipients) and identified 106 successful colonization events, 109 failed colonization events, and 700 ambiguous colonization events (Supplementary Table 6). Our stringent criteria classified the vast majority of all genome/recipient pairs as ambiguous colonization events. Nevertheless, due to the relatively large number of donor MAGs and FMT recipients in our study, we were left with 352 MAG/recipient pairs with unambiguous colonization phenotypes for downstream analyses.

Using colonization information from our improved model that took into consideration the presence and absence of distinct subpopulations and their origins (Supplementary Figure 4), we then tested if colonization success was correlated with population fitness or population dose, which we define here as the relative abundance of a given population in the transplanted donor stool sample (we calculated the relative abundance of individual populations using the second and third quartile of the mean coverage of populations in metagenomes, a practice that reduces the impact of variation in coverage as a function of hypervariable genomic islands and/or conserved genomic regions that recruit reads from multiple populations). For Donor A populations, colonization outcome was significantly correlated with both dose (Wald test, AUC=0.73, p=7.7e-05) and fitness (Wald test, AUC=0.76, p=6.3e-06) (Figure SIb,c). But combining both measures as predictive variables did not substantially improve the performance of our colonization model (AUC=0.82) (Figure SIc). This was likely due to the small, but significant, correlation between dose and fitness in Donor A MAGs ($R^2$=0.053, p=0.0070) (Figure SId). When the fitness of a microbial population is reflected in its relative abundance, the effect of fitness on colonization outcome may be masked by an apparent dose effect. In contrast to Donor A, the fitness of Donor B populations and their relative abundance in Donor B samples were not correlated ($R^2$=0.0012, p=0.61) (Figure SId), providing us with an ideal case to analyze these two factors independently. Indeed, there was no correlation between dose of a microbial population in Donor B transplant samples and colonization outcome in recipients post-FMT (Wald test, AUC=0.56, p=0.09). Instead, we found a significant correlation between the fitness of each population and the colonization outcome (Wald test, AUC=0.70, p=9.0e-07) (Figure SIc).

Taken together, our findings suggest that fitness of a microbial population as measured by its prevalence across global gut metagenomes can predict its colonization success better than its abundance in the donor stool sample, giving credence to the role of adaptive rather than neutral ecological processes in colonization. This finding contrasts with previous studies which suggested that the abundance of a given population in the donor sample was an important determinant of colonization (Podlesny and Florian Fricke, 2020; Smillie et al., 2018). However, these analyses included many recipient samples collected less than one week after FMT and it is likely that their observations were influenced by the presence of transient populations. Indeed, samples collected immediately after FMT are more likely to inflate the number of colonization events, whereas longitudinal sampling over a longer time course can distinguish transient populations from those that successfully colonized the recipients. We cannot definitively test this hypothesis as most of our recipient samples were collected at least one week after FMT. Still, on average 12% of the donor populations detected in our recipients a week after FMT were no longer detected after a month (Figure 1, Supplementary Table 3). Overall, our stringent criteria to determine colonization outcome and the extended post-FMT sampling period likely enabled us to study the long-term engraftment of successful and potentially low-abundance colonizers, instead of high-abundance transient populations that may be dominant directly after FMT.

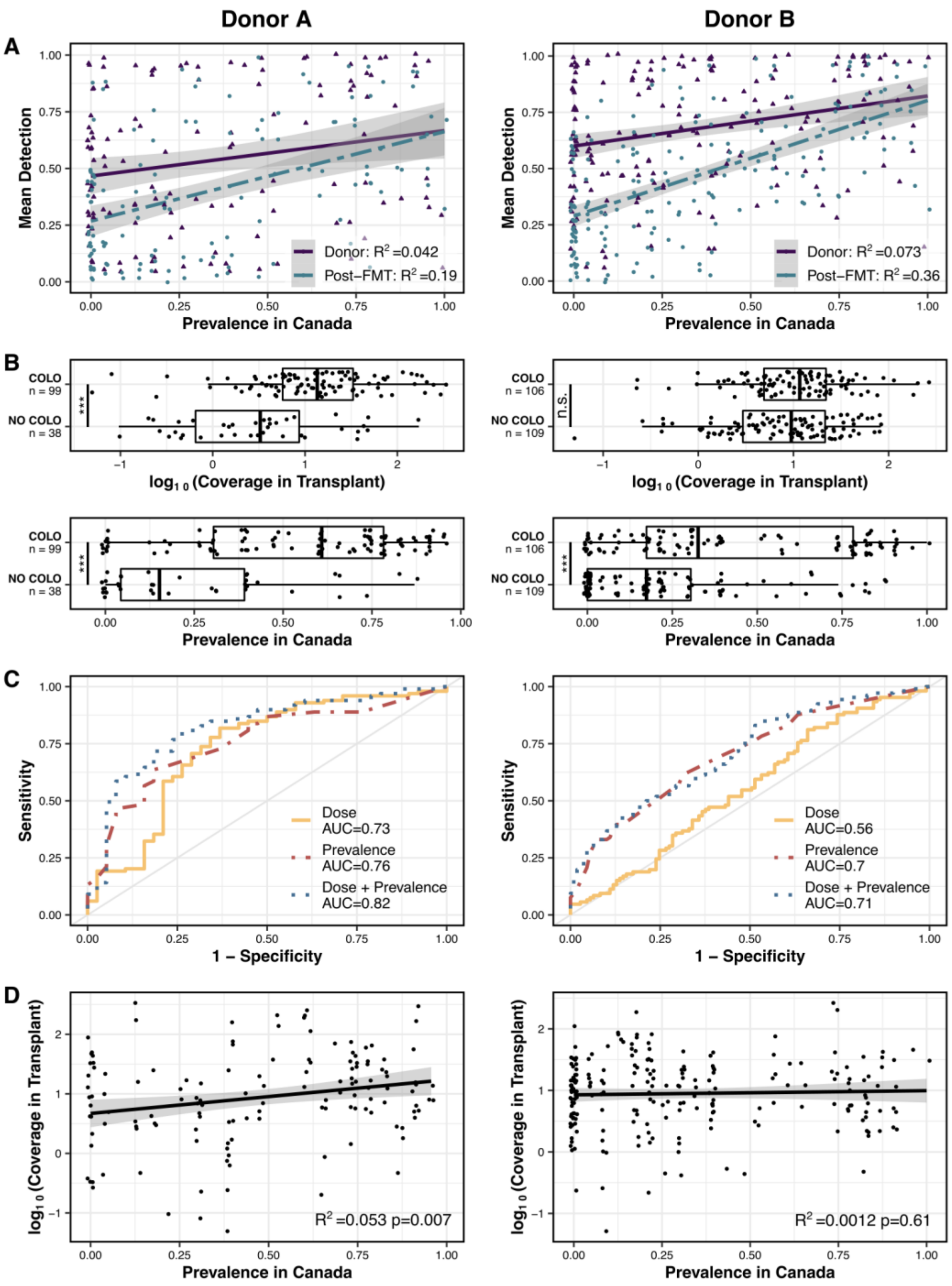


**Figure SI. Relationships between dose, prevalence, and colonization outcome.** Left: Donor A. Right: Donor B. (a) Linear regression models of mean detection of each MAG in either donor or recipient post-FMT samples as a function of prevalence. (b) Colonization outcome of MAG/recipient pairs as a function of MAG dose or MAG prevalence. Significance calculated by Wald test. (c) Receiver operator curves (ROCs) for logistic regression models of colonization. (d) Linear regression models of dose as a function of prevalence. The grey shaded areas in panels (a) and (d) represent the 95% confidence intervals. To examine the association between dose and/or prevalence with colonization outcome in this regression analysis shown in the figure, we built binomial logistic regression models using the R stats `glm` function. We used the R stats `predict` function and the R pROC `roc` function to evaluate our models by creating receiver operating characteristic (ROC) curves and calculating the area under the ROC curve (AUC). To determine the correlation between dose and prevalence, we performed linear regression using the R stats `lm` function. We used the R tidyverse package, including ggplot2, to visualize boxplots, scatterplots, and ROC curves. The URL https://doi.org/10.6084/m9.figshare.15138720 serves a high-resolution version of this figure.

Accurately distinguishing the role of dose versus fitness in colonization success is further compounded by the fact that microbial populations that are prevalent across human populations may also tend to be more abundant. This is well illustrated by Donor A. Fortunately, the abundant populations in Donor B did not reflect prevalent microbes in healthy adult guts, which demonstrated the importance of fitness as a determinant of colonization success compared to dose without the confounding effect of a correlation between fitness and dose. Thus, it is a theoretical possibility that colonization success is purely driven by adaptive forces and is not influenced by dose, at all. However, while our data assign a larger role to adaptive forces with confidence, a more accurate determination of the proportional influence of adaptive versus neutral processes in colonization requires a much larger dataset.

### References

Denef VJ. 2019. Peering into the Genetic Makeup of Natural Microbial Populations Using Metagenomics In: Polz MF, Rajora OP, editors. Population Genomics: Microorganisms. Cham: Springer International Publishing. pp. 49–75.

Maignien L, DeForce EA, Chafee ME, Eren AM, Simmons SL. 2014. Ecological succession and stochastic variation in the assembly of Arabidopsis thaliana phyllosphere communities. *MBio* **5**:e00682–13.

Podlesny D, Florian Fricke W. 2020. Microbial Strain Engraftment, Persistence and Replacement after Fecal Microbiota Transplantation. *medRxiv* 2020.09.29.20203638.

Quince C, Delmont TO, Raguideau S, Alneberg J, Darling AE, Collins G, Eren AM. 2017. DESMAN: a new tool for de novo extraction of strains from metagenomes. *Genome Biol* **18**:181.

Smillie CS, Sauk J, Gevers D, Friedman J, Sung J, Youngster I, Hohmann EL, Staley C, Khoruts A, Sadowsky MJ, Allegretti JR, Smith MB, Xavier RJ, Alm EJ. 2018. Strain Tracking Reveals the Determinants of Bacterial Engraftment in the Human Gut Following Fecal Microbiota Transplantation. *Cell Host Microbe* **23**:229–240.e5.
